# Alternative Macrophage Activation Requires Granulocyte-Derived Lipids

**DOI:** 10.1101/2024.02.17.580795

**Authors:** Tatyana Veremeyko, Anders Vik, Trond Vidar Hansen, Eugene D. Ponomarev

## Abstract

Lipid metabolism is important for alternative (M2) macrophage activation, but it remains unclear whether it is essential for this process, and the underlying mechanisms have yet to be determined. It is also unclear how the M2 phenotype is induced during the resolution of inflammation and tissue repair. We used large-scale RNA-sequencing and high-throughput lipidomics approaches to identify that during blood coagulation, granulocytes release certain lipids that are loaded onto high-density lipoproteins (HDLs), which then act via scavenger receptors SR-B1 and SR-B3 - to mediate the expression of 26 of 32 M2-associated genes and oxidative phosphorylation (OXPHOS) in macrophages. We found that in the absence of these lipids, interleukin-4 (IL-4) induced only 6 of 32 M2 markers, and this cytokine alone failed to induce OXPHOS. We also found that the HDL-associated C18:0, C18:2, and C20:3 fatty acids (FAs) were the lipids that most potently induced the M1-to-M2 transition. Furthermore, we determined that during IL-4-mediated inflammation, the inhibition of blood coagulation, or platelet and granulocyte depletion, results in a substantial decrease in M2 marker expression. Thus, this study identified that granulocyte-derived FAs mediate the reprogramming of macrophages towards the M2 phenotype, thereby bridging the processes of blood coagulation, inflammation, and repair.

**Highlights:**

- Activation of macrophages with IL-4 or IL-13 requires indispensable cofactor
- The cofactor consists of high-density lipoproteins (HDLs) complexed with lipids
- During blood coagulation, granulocytes produce lipids that are loaded on HDLs
- HDL-associated fatty acids were the most potent lipids to induce M2 phenotype

**Graphic Abstract:** 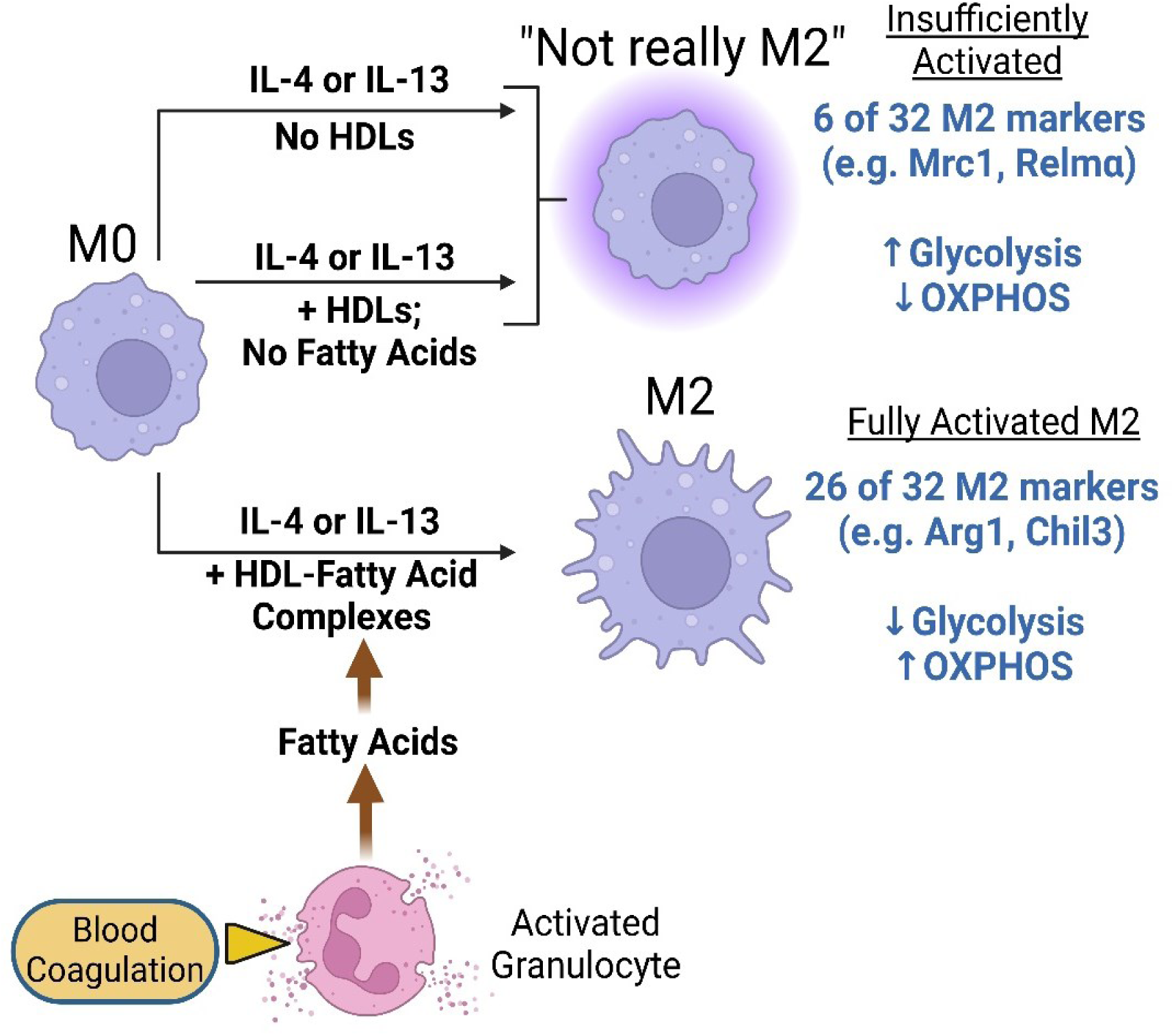

## Introduction

Immunometabolism is a new research direction in inflammation biology that is aimed at finding pathways via which immune cells can be programmed to shift from a pathological phenotype towards a ‘healing’ phenotype that mediates tissue repair and restoration of homeostasis. ^1^ For example, several studies have focused on the metabolic reprogramming of proinflammatory M1 macrophages towards alternatively activated M2 macrophages that promote the resolution of inflammation, wound healing, and tissue remodeling. ^2–4^ Lipid metabolism is important for M2 activation, ^5^ whereas glycolysis is typically used by M1 macrophages. ^6^ Specifically, M2 macrophage oxidative phosphorylation (OXPHOS) has been shown to involve fatty acid (FA) oxidation. ^3,7–11^

FA oxidation has also been found to be unnecessary for M2 activation, and its role remains controversial. ^11–13^ It was demonstrated that FA-containing triglycerides (TGs) loaded onto low-density lipoproteins (LDLs) are delivered to macrophages via scavenger receptor class B member 3/cluster of differentiation 36 (SR-B3/CD36) receptors. ^9^ However, the aforementioned study did not determine the main source of FAs during inflammation and whether it was an external source or an internal source.

In the current study, we found that M2 activation requires extrinsic serum-derived free FAs as a cofactor for IL-4 or IL-13. This cofactor consists of high-density lipoproteins (HDLs) loaded with saturated and unsaturated FAs, and the loading process occurs during blood coagulation and IL-4/IL-13-mediated (type 2) inflammation. Most importantly, our study indicates that granulocytes provide an external pool of FAs during blood coagulation by performing ‘targeted feeding’ of macrophages, thereby mediating the resolution of inflammation and tissue repair by promoting the M1-to-M2 transition.

## Results

### IL-4 or IL-13 requires a previously unknown serum cofactor

We previously observed that the batch of fetal bovine serum (FBS) used for cell culture affected the level of mRNA expression of main M2 marker Arginase 1 (*Arg1*) in macrophages, which is consistent with another earlier observation that FBS may have pro-M2 and anti-M1 activities.^14^ The variation between batches of FBS is attributable to its being collected at different stages of fetal development or contaminated with adult bovine serum (ABS). Macrophages can survive in serum-free media for more than 24 h ^14^, and their viability does not significantly decrease over four days in culture (data not shown). We examined the role of serum factors in *Arg1* expression by incubating murine bone-marrow-derived macrophages (BMDMs) in serum-free media (untreated); serum-free media with IL-4; or in media with IL-4 supplemented with FBS, or ABS, or adult human serum (AHS), or adult mouse serum (AMS). We found that incubation of murine BMDMs in IL-4 alone for 24 h failed to induce *Arg1* expression, but that incubation of murine BMDMs in IL-4 with FBS, ABS, AHS, and AMS induced *Arg1* expression (**Figure 1A**). The effect was stronger in the case of adult serums vs. FBS. In addition, *Arg1* (mRNA) and ARG1 (protein) concentrations peaked on days 4 and 8, respectively, in the presence of FBS or ABS, with a much lower concentration of ARG1 induced by FBS than by ABS (**Figure 1B** and **Figure S1A**, **Supplemental Material**). IL-4 alone failed to induce *Arg1* and ARG1 until day 4 (when the cells remained viable in a serum-free medium), and arginase enzymatic activity required the presence of FBS or ABS (**Figure 1B**). Fractionation experiments (**Figure 1C**) further demonstrated that a previously unknown serum cofactor for IL-4 (**Figure 1D–G**) and IL-13 (**Figure S1B–D**, **Supplemental Material**) had a molecular weight of more than 100 kDa (>100 kDa) in both FBS and ABS (‘FBS-hi’ and ‘ABS-hi’ fractions, respectively; **Figure 1C**), and substantially upregulated mRNA for Arg1 and chitinase-like 3 (*Arg1* and *Chil3*, **Figure 1D**), proteins for ARG1 and CHIL3 (**Figure 1E, F**), and arginase enzymatic activity (**Figure 1G** and **Figure S1D**, **Supplemental Material**), but downregulated mRNA for nitric oxide synthase 2 (*NOS2*), NOS2 protein, and mRNA for tumor necrosis factor (*TNF*) in interferon-gamma/lipopolysaccharide-activated M1 macrophages (**Figure S1E–H**). The experiments depicted in **Figure 1D–G** and the following experiments (e.g., those depicted in Figure 2) were performed in the presence of the ‘FBS-lo’ (<100 kDa) fraction that contained most of the growth factors and regulatory cytokines, such as all isoforms transforming growth factor beta (TGFβ), that could affect M2 activation, ^14^ cell viability, and proliferation. ^15^

**Figure 1.**
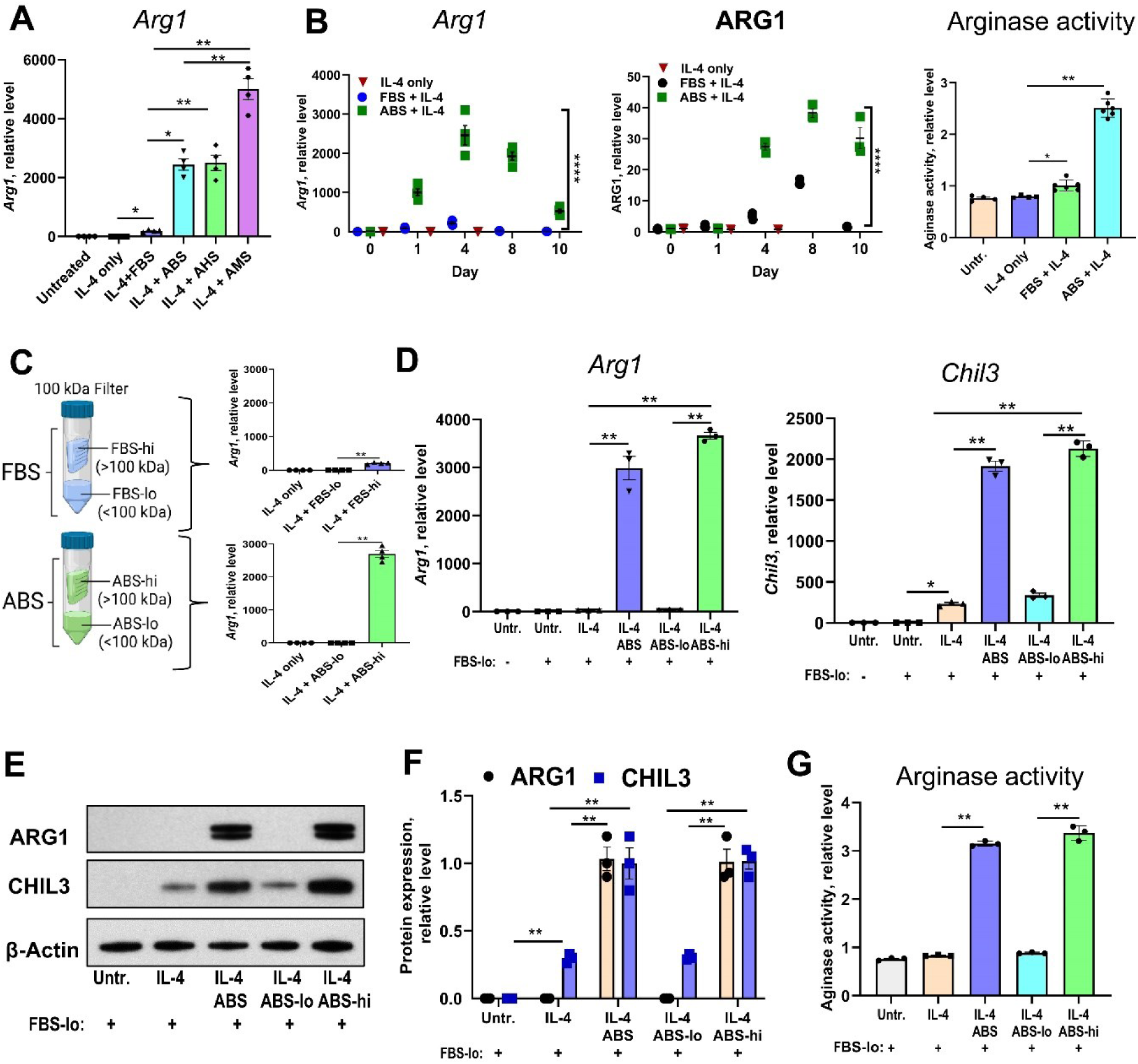
The influence of serum cofactor for interleukin 4 on the expression of M2 markers. (**A**) Species-specific synergistic effect of fetal bovine serum vs. adult bovine, adult human, and adult mouse serums on the expression of mRNA for Arginase 1 (Arg1) in interleukin - 4 (IL-4) - treated mouse bone marrow-derived macrophages (BMDM). (**B**) Comparison of effects of fetal bovine serum (FBS) and adult bovine serum (ABS) on the kinetics (day 1, 4, 8, and 10) of expression of Arginase 1 on mRNA (*Arg1*, left panel, real-time RT-PCR) and protein (ARG1, middle panel; quantitative western blotting) levels, and arginase enzymatic activity (right panel) in IL-4 treated or untreated murine BMDM (see Methods and **Figure S1A**, **Supplemental Material**). (**C**) Identification of more than 100 kDa (>100 kDa) high molecular weight fractions of FBS and ABS (FBS-hi and ABS-hi, respectively) as an active component for cofactor for IL-4 to induce *Arg1* in BMDM. (**D-G**) Comparisons of effects ABS, ABS-lo (<100 kDa), and ABS-hi (>100 kDa) serum fractions on the expression of M2 markers Arg1 and Chitinase-like protein 3 (Chil3) on mRNA (**D**) and protein (**E, F**) levels, as well as arginase enzymatic activity (**G**) in IL-4 - treated or untreated BMDM. For (**A-D**), (**F**), (**G**): *p<0.05, **, p<0.01, ****, p<0.0001, data represented as mean ± s.e.m.; see **Table S1** for detailed statistics. <u>Abbreviations</u>: ABS, adult bovine serum; AHS, adult human serum; AMS, adult mouse serum; FBS, fetal bovine serum; FBS-lo, low molecular weight (>100 kDa) fraction of FBS; Untr., untreated. See also **Table S1** and **Figure S1, Supplemental Material**.

**Figure 2.**
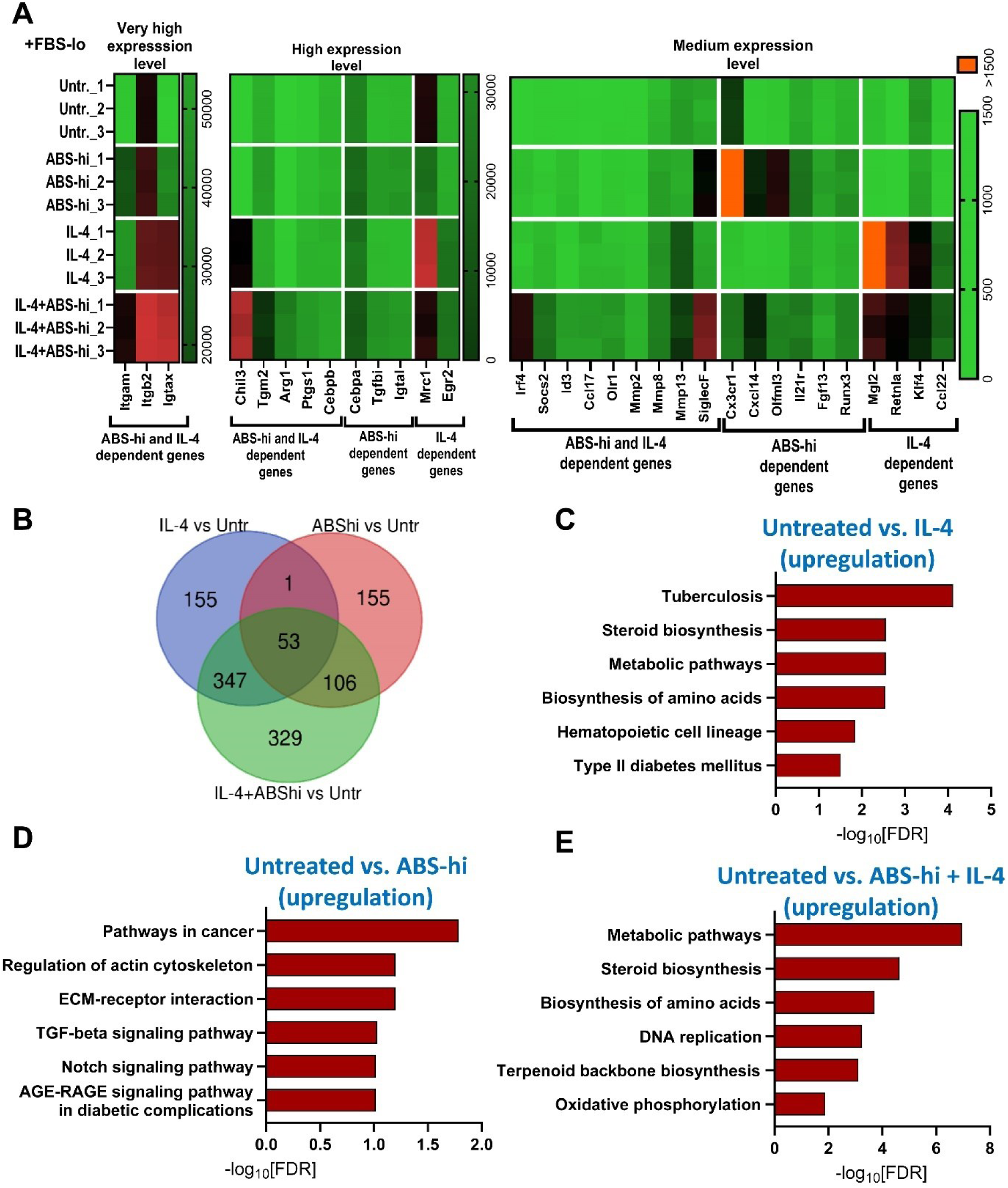
Transcriptomic analysis of the expression of M2-associated genes and signaling pathways in IL-4-stimulated macrophages in the presence of ABS-hi cofactor. (**A**) A hit map of expression of M2-associated genes in untreated murine BMDM, vs. macrophages treated with ABS-hi, or IL-4, or IL-4 and ABS-hi (IL-4 + ABS-hi) is shown for RNAseq assay (n=3). (**B**) Venn diagram of a total number of upregulated genes in untreated mouse BMDM vs. macrophages treated with IL-4 (left), ABS-hi (right), or IL-4 + ABS-hi (bottom). (**C-E**) KEGG analysis of specific pathways that were activated (upregulated) in macrophages treated with IL-4 alone (**C**), ABS-hi alone (**D**), or both IL-4 with ABS-hi (**E**) indicated the importance of the presence of both ABS-hi and IL-4 for upregulation of oxidative phosphorylation and other metabolic pathways. <u>Abbreviations</u>: ABS, adult bovine serum; ABS-hi, high molecular weight (>100 kDa) fraction of ABS; FBS-lo, low molecular weight (<100 kDa) fraction of FBS; FDR, False Discovery Rate; Untr, untreated. See also **Table S1** and **Figure S2, Supplemental Material.**

Our further transcriptomics analysis revealed that more than 80% of IL-4-upregulated M2-related genes require the presence of the previously unknown ABS-hi serum cofactor (**Figure 2A, B**), except for several M2-associated genes (mannose receptor C-type 1 (*Mrc1*), early growth response 2 (*Egr2*), macrophage galactose *N*-acetyl-galactosamine specific lectin 2 (*Mgl2*), and resistin-like alpha (*Retnla*)), which are induced by IL-4 alone without the presence of ABS-hi cofactor (**Figure 2A**, and **Figure S2A, B, Supplemental Material**).

Kyoto Encyclopedia of Genes and Genomes (KEGG) analysis demonstrated that the previously unknown ABS-hi serum cofactor substantially enhances IL-4 activity via the upregulation of genes involved in multiple metabolic pathways, including the biosynthesis of amino acids, steroids, and OXPHOS, whereas ABS-hi alone upregulates the phosphoinositide 3-kinase (PI3K)/Akt pathway, which is involved in cancer, as well as transporters of amino acids and fatty acids (**Figure 2C-E**). Thus, more than 80% of IL-4-upregulated M2-associated genes and downstream metabolic pathways require the previously unknown ABS-hi serum cofactor.

### Coagulation activates the previously unknown serum cofactor

We found that ABS was more efficient than FBS in inducing *Arg1* and ARG1 expressions and its enzymatic activity, suggesting that the blood plasma composition and cellular subsets in adult vs. fetal blood may affect this process. Thus, we hypothesized that blood coagulation influences the activity of the previously unknown serum cofactor required for generating the M2 phenotype in murine BMDMs. We tested this hypothesis by comparing the ability of ABS and adult citrate bovine plasma (CBP) to induce *Arg1* expression in macrophages and found that in combination with IL-4, ABS but not CBP induced *Arg1* expression (**Figure S2C**, **Supplemental Material**). Since in the case of ABS vs. CBP we need to compare products from different animals, we were performing experiments using adult mouse citrate plasma (MCP) and AMS, respectively. For AMS and CMP preparations, the blood was collected from age- and sex-matched groups of syngeneic animals, and we used an additional control, namely the addition of sodium citrate (SC) to sera (AMS + SC) immediately before experiments (**Figure 3A**). We found that CMP had no, or very weak effect compared to AMS for the induction of expression of M2 markers, which affected the expressions of Arg1 and Chil3 on mRNA and protein levels (**Figure 3B, D**), and arginase enzymatic activity (**Figure 3E**). Given that we obtained the same results when we used heparin (H) instead of SC as an anticoagulant (**Figure S2D-G**, **Supplemental Material**), we concluded that coagulation is required for the functional activity of the previously unknown serum cofactor.

**Figure 3.**
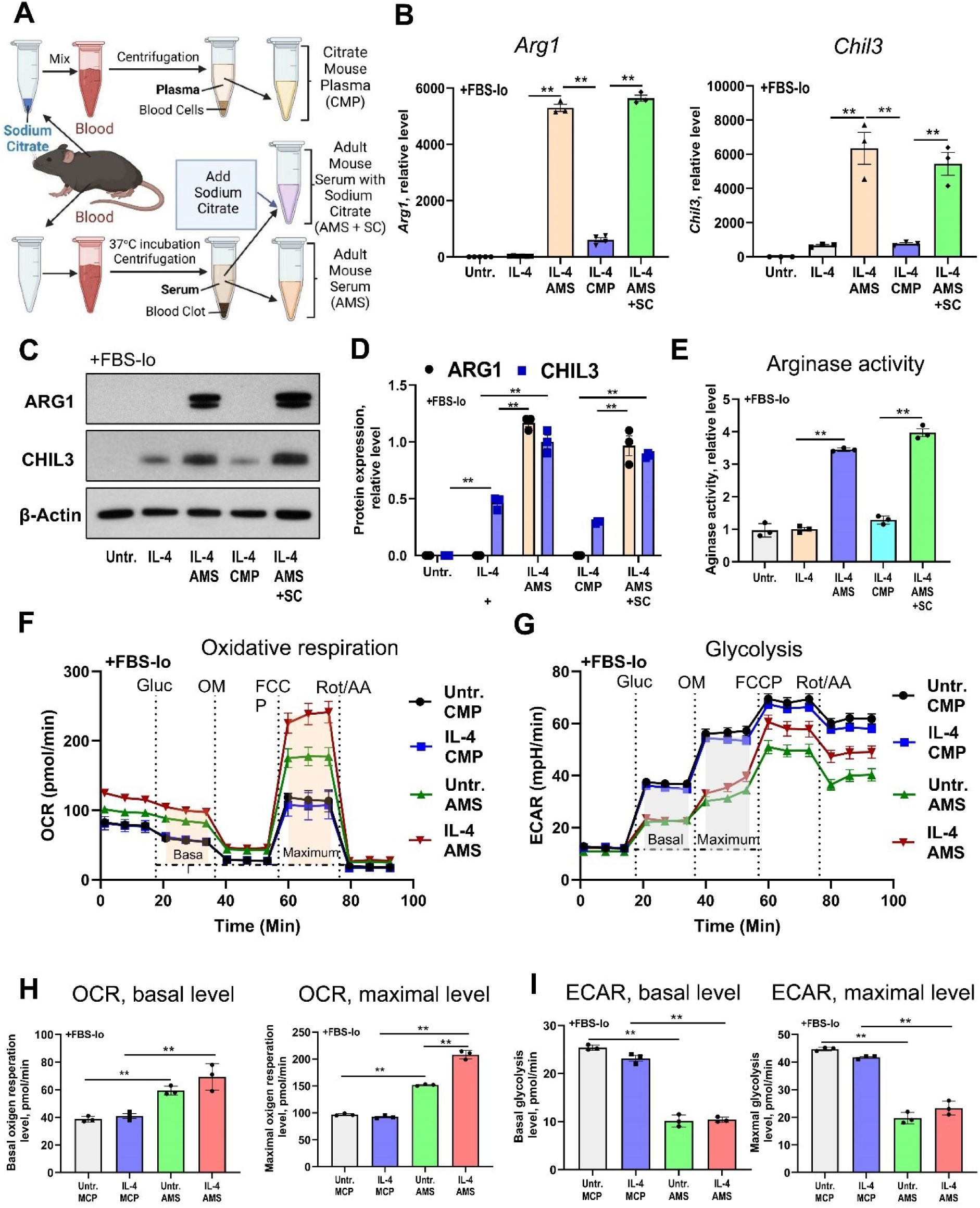
The influence of the blood coagulation on the functional activity of cofactor for IL-4 to induce the expression of M2 markers and oxidative phosphorylation in macrophages. (**A**) Schematic diagram for ex-vivo preparations of adult mouse serum (AMS), citrate mouse plasma (CMP), and AMS with subsequently added sodium citrate (AMS + SC) substances (see Methods). (**B-E**) Comparisons of effects of AMS, CMP, and AMS+SC substances on the expressions of M2 markers Arg1 and Chil3 on mRNA (**B**) and protein (**C, D**) levels and arginase enzymatic activity (E) in IL-4-treated or untreated murine BMDM (see Methods). (**F-I**) Comparison of effects of AMS and CMP on oxidative respiration (**F, H**) and glycolysis (**G, I**) rates in IL-4-treated or untreated murine BMDM (see Methods). Representative real-time kinetics of oxygen consumption (**F**) and extracellular acidification (**G**) rates (OCR and ECAR, respectively) are shown in (**F, G**). Quantifications of basal (left panel) and maximal (right panel) levels of OCR (**H**) and ECAR (**I**) are shown in (**H, I**). For (**B**), (**D**), (**E**), (**H**), (**I**): **, p<0.01; data represented as mean ± s.e.m.; see **Table S1** for detailed statistics. <u>Abbreviations</u>: AMS, adult mouse serum; AA, antimycin A, CMP, citrate mouse plasma; ECAR, extracellular acidification rate; FBS-lo, low molecular weight (>100 kDa) fraction of FBS; FCCP, Carbonyl cyanide-4 (trifluoromethoxy) phenylhydrazone; Gluc, glucose; OCR, oxygen consumption rate; R, rotenone; SC, sodium citrate; OM, oligomycin; Untr., untreated. See also **Table S1** and **Figure S2, Supplemental Material**.

### IL-4 alone cannot induce OXPHOS

IL-4 induces OXPHOS in mouse macrophages, ^8,9^ and thus we tested whether a previously unidentified serum cofactor in AMS or ABS was required for the stimulation of OXPHOS, as was suggested by our KEGG pathway analysis (**Figure 2E**). We used citrate mouse plasma (CMP) as a negative control, as it does not induce ARG1 expression in macrophages (**Figure 3B**), and also supplemented macrophages with an FBS-lo fraction that contained most of the required growth factors. We found that IL-4 with CMP failed to increase the basal or maximal oxidative respiration levels above those in unstimulated murine BMDMs incubated with CMP (**Figure 3F, H**). In addition, AMS alone significantly stimulated oxidative respiration in macrophages (i.e., AMS accounted for more than 50% of the observed stimulation), and stimulation was further enhanced by IL-4 **(Figure 3F, H**). Interestingly, however, AMS alone substantially decreased glycolysis, but IL-4 and CMP did not significantly affect the basal or maximal levels of glycolysis (**Figure 3G, I**). Thus, the previously unknown serum cofactor alone, or in combination with IL-4, increased OXPHOS and decreased glycolysis in murine BMDMs.

### The serum cofactor consists of lipoproteins and lipids

Next, we aimed to identify the previously unknown serum cofactor in the ABS-hi fraction using the multistep fractionation strategy shown in **Figure S3A** (**Supplemental Material**). We tested the biological activity of each fraction using the murine macrophage line RAW264.7 as a test system to measure *Arg1* expression, which was normalized as a percentage of its level of expression at the previous step of fractionation (**Figure S3B-D**). We found that the previously unknown serum cofactor was absorbed into the hydrophobic column, and subsequent elution indicated the presence of the highly hydrophobic protein apolipoprotein A1 (APOA1) eluted at Steps 5 and 6 (**Figure S3E**) and other less hydrophobic proteins, namely bovine serum albumin (BSA), immunoglobulins, and complement eluted at Step 5 (**Figure S4A, B**, **Supplemental Material**). Bands 1 and 2 obtained in Step 5 contained C3 a major component (**Figure S4B**), and APOA4 (Band 1) and APOA1 (Band 2) as minor components (not shown), which are both components of HDLs and were further enriched after Step 6 (**Figure S3E**). All of the less hydrophobic proteins examined did not affect *Arg1* expression, as demonstrated by tests using BSA and immunoglobulin preparations (**Figure S4C**), and by comparing sera from adult wild-type (WT), C3^−/−^ and C4^−/−^ mice (**Figure S4D**), and normal sera from adult humans and human adult sera with depleted complement components C3, C4, or both C3 and C4 (not shown). APOA1 is a main structural component of HDLs, and its presence in the most active fraction after Step 6 indicated that the unknown serum cofactor contained HDLs and most likely was complexed with lipids, as pure APOA1 that was eluted completely from the hydrophobic column by extensive volumes of 8M urea did not exhibit substantial biological activity (data not shown).

Subsequently, we verified that the previously unknown serum cofactor contains lipoproteins complexed with lipids by using gradient ultracentrifugation to isolate HDLs, LDLs, and all other protein fractions from ABS and AMS (**Figure 4A**). In both types of sera, HDLs were more active than LDLs and non-lipoprotein ‘protein’ fractions (**Figure 4B**). In ABS, LDLs also demonstrated *Arg1*-inducing activity, whereas, in AMS, they exhibited *Arg1*-inducing activity that was not significantly greater than that exhibited by a protein fraction or negative control (**Figure 4B**, ABS vs. AMS). Moreover, the treatment of HDLs in AMS with proteases and glycosidases (**Figure 4C**) did not significantly decrease their *Arg1*-inducing activity in RAW264.7 cells, indicating that HDL-associated lipids, but not proteins or carbohydrates, are important for this activity (**Figure 4D**). Finally, we isolated lipids from AMS and created artificial liposomes (**Figure 4C**) that exhibited ∼40% of the ARG1-inducing activity of HDLs (**Figure 4D**, Liposomes), which revealed that lipoprotein-associated lipids are key cofactors of IL-4.

**Figure 4.**
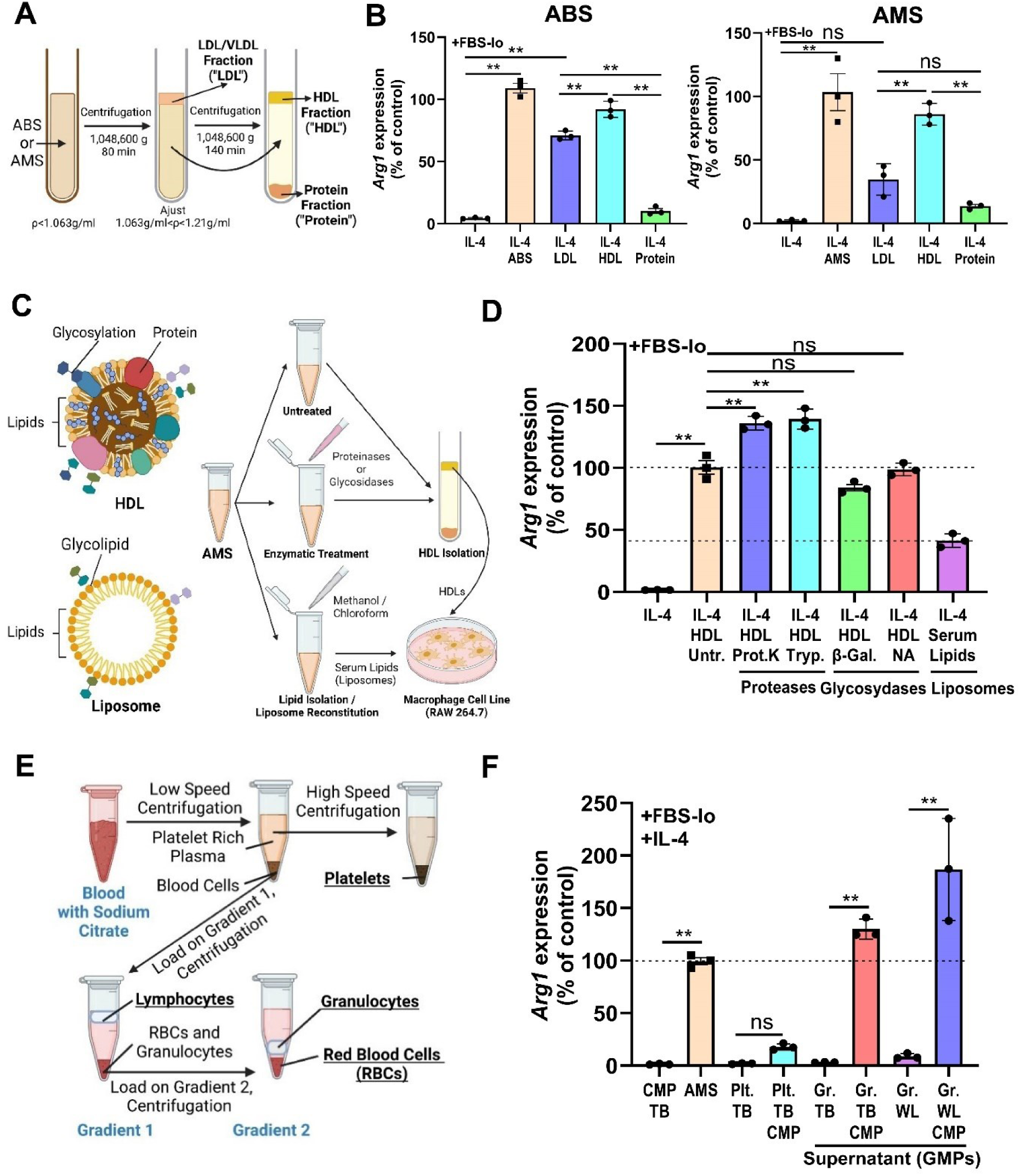
Analysis of low- and high-density lipoproteins and populations of blood cells as sources of lipids that potentiate M2 activation. (**A**) Schematic diagram of isolation LDL, HDL, and non-lipoprotein ‘protein’ fractions from adult murine or bovine serums. (**B**) Comparisons of effects of unfractionated ABS or AMS vs. LDLs, HDLs, and non-lipoprotein ‘protein’ fractions on *Arg1* expression in IL-4-treated macrophage cell line (see Methods). (**C**) Schematic images of structures of HDLs and artificial liposomes, and the experimental design for enzymatic treatment of murine HDLs from AMS with proteases and glycosidases, as well as plan for isolation of mouse serum lipids and the creation of artificial liposomes (see Methods). (**D**) Effects of enzymatic treatments of mouse HDLs with proteinases (proteinase K and trypsin), glycosidases (galactosidase and neuraminidase), addition of artificial liposomes containing AMS-derived lipids on *Arg1* expression in IL-4-treated macrophages. (**E**) Experimental design of isolation of platelets, mononuclear leukocytes (‘lymphocytes’), granulocytes, and erythrocytes (red blood cells, RBCs) by separation of platelet-rich plasma by low-speed centrifugation followed by two-step Percoll density gradient centrifugation. (**F**) Blood-derived cell populations were activated with thrombin (TB) or lysed with water to make water lysate (WL), their supernatants were mixed with CMP (as a source of lipoproteins) and then added to macrophages for analysis of *Arg1* expression (refer to **Figure S5A-C** and Methods). Supernatants that contained granulocyte-derived microparticles (GMPs) or WL, which were mixed with CMP (but not without CMP), exhibited significant upregulation of *Arg1*. For (**B**), (**D**), (**F**): **, p<0.01; ns, not significant; data represented as mean ± s.e.m.; see **Table S1** for statistics). <u>Abbreviations</u>: β-Gal., β-galactosidase; Gr., granulocytes; GMPs, granulocyte-derived microparticles; NA, neuraminidase; Plt., platelets; Prot.K, proteinase K; Tryp., trypsin; TB, thrombin; WL, water lysate; Untr., untreated. See also **Table S1** and **Figure S5, Supplemental Material.**

### Granulocytes are the main source of lipids

It is well established that, in addition to platelets, activated granulocytes (mostly neutrophils) also contribute to blood coagulation and clot formation *in vitro* and *in vivo* via various mechanisms, including the release of granule substances, DNA (as part of ‘neutrophil extracellular traps’), and microparticles. ^16,17^ Accordingly, we hypothesized that platelets and granulocytes are the main sources of lipids that become associated with HDLs during blood coagulation. We tested this hypothesis by performing a multistep separation of platelets, granulocytes, erythrocytes (red blood cells, RBCs), and blood mononuclear leukocytes (i.e., ‘lymphocytes’, as these cells were mostly T and B cells, see Methods) as shown in **Figure 4E** and **Figure S5A** (**Supplemental Material**). The purities of the isolated populations were assessed by flow cytometry (**Figure S5B**, **Supplemental Material** and Methods). We then activated cells in each population with thrombin (TB), a potent coagulator that directly activates platelets, and directly or indirectly^18^ (possibly via platelets aggregated with granulocytes ^19^) induces granulocyte degranulation, ^20^ and could also potentially activate blood mononuclear leukocytes via protease-activated receptors. ^21^ As a control for the passive release of granules and cytoplasmic substances, we lysed cell subsets with water to obtain water lysates (WL). After the activation of all cell populations with TB or after making WL, we used low-speed centrifugation to collect supernatants containing microparticles or cytoplasm substances from granulocytes, mononuclear lymphocytes, and RBCs, but not platelets, as they are too small (2-3 µl) to separate their microparticles from cells by low-speed centrifugation (see Methods). We found that supernatants from TB-activated granulocytes and those from WLs mixed with CMP (the source of lipoproteins such as HDLs) upregulated *Arg1* expression by 125%–175% of the level by which it was upregulated by AMS (**Figure 4F**). In contrast, TB-activated platelets combined with CMP upregulated *Arg1* expression to a level 15%–20% of the level by which it was upregulated by AMS, although the difference between these two levels of upregulation was not statistically significant (**Figure 4F**). Similarly, RBCs and lymphocytes did not upregulate the expression of *Arg1* to a statistically significant extent (**Figure S5C**, **Supplemental Material**). Moreover, microparticles that were not combined with CMP and had been isolated from TB-activated platelets and granulocytes failed to regulate *Arg1* expression, confirming the critical role of lipoproteins in this process (**Figure 4F** and **Figure S5C**).

Next, due to the limited amount of circulating granulocytes present in mice ^22^, we isolated large amounts of a greater than 95% pure population of CD11b^+^Ly6G^+^ granulocytes from murine bone marrow (**Figure S5D, E** and Methods) and found that they did not differ from blood-derived granulocytes in their ability to induce expression of *Arg1* (**Figure S5F**). Subsequently, we performed flow cytometry of granulocyte-derived microparticles (GMPs, see Methods and **Figure S5G**) that are known to contain lipids such as TGs ^23^ and found that stimulation of granulocytes with TB caused them to secrete GMPs and TGs. The secretion of GMPs (**Figure 5A-D**) and TGs (**Figure 5A, C, E**) was significantly decreased by pre-treating cells with two neutral sphingomyelinase (nSMase) inhibitors (**Figure 5A**) that block the release of microparticles and exosomes from many types of cells. ^24–26^ Supernatants from inhibitor-pretreated granulocytes activated with TB failed to induce the expression of *Arg1* in macrophages (**Figure 5F**), indicating the importance of GMP production by granulocytes to induce the expression of Arg1 in the macrophage cell line. Finally, *in vivo* platelet or granulocyte depletion, or intravenous injections of nSMase inhibitors, rendered subsequently collected AMS inactive, but the addition of these inhibitors to prepared AMS did not affect *Arg1* expression in macrophages (**Figure 5G**). Overall, these aforementioned data indicate that granulocytes serve as a main source of lipids for circulating lipoproteins that induce M2 activation.

**Figure 5.**
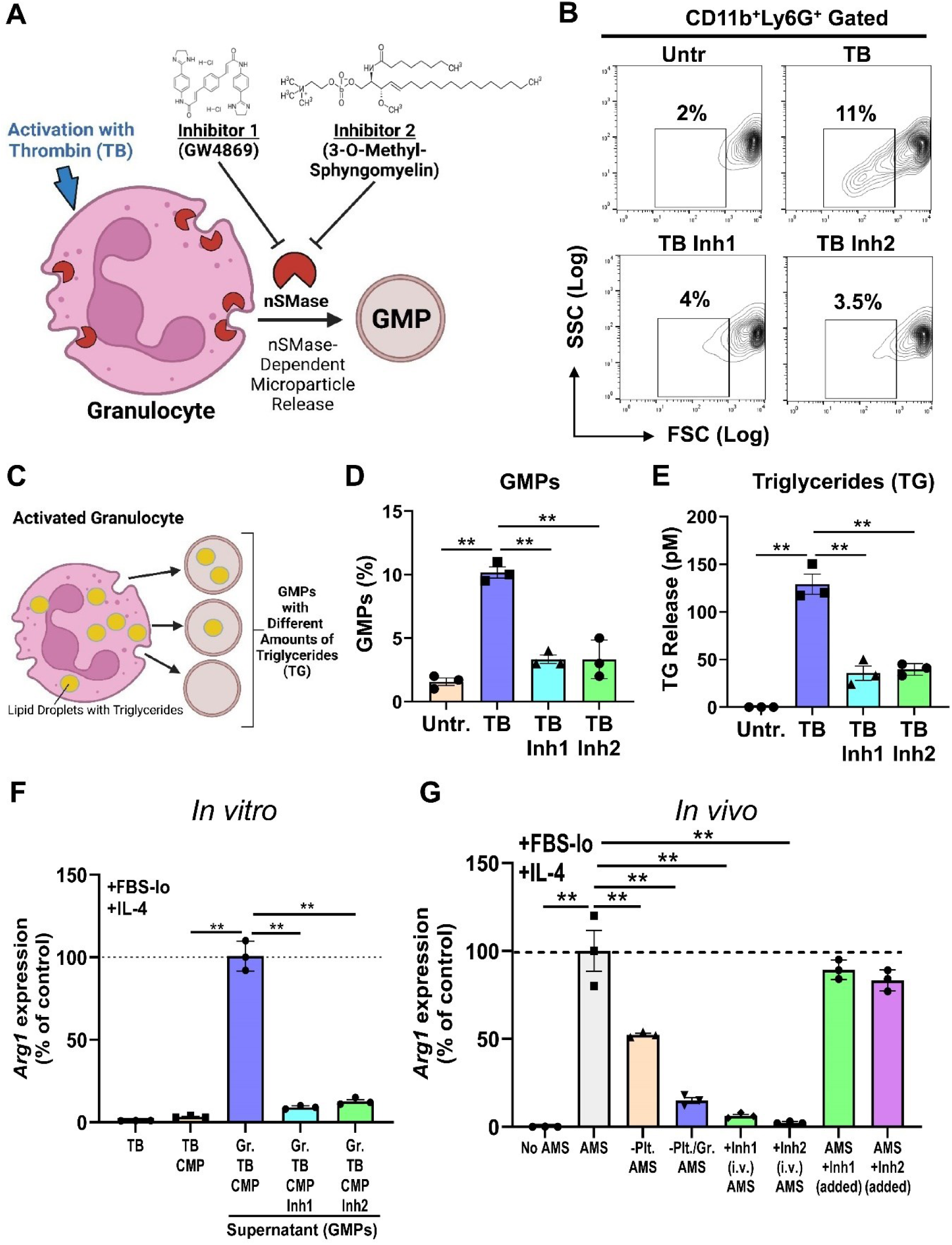
Analysis of the role of granulocytes and their secretory products as the main source of lipids that potentiate M2 activation. (**A**) Schematic diagram of inhibition of secretion of granulocyte-derived microparticles (GMPs) by granulocytes activated with thrombin (TB) using two inhibitors (‘Inhibitor 1’ and ‘Inhibitor 2’) of neutral sphingomyelinase (nSMase, see Methods). (**B**) Flow cytometry analysis of percentages of GMPs in untreated granulocytes or the cells pre-treated with two nSMase inhibitors to prevent the formation of GMPs (Inh1 and Inh2, see also **Figure S5D-G, Supplemental Material** and Methods). (**C**) Schematic image of secretion of GMPs by activated granulocytes, which contain various amounts of lipid droplets with triglycerides (TG). (**D, E**) Quantitative analysis of GMP (**D**) and triglyceride (**E**) releases in untreated granulocytes or the cells pre-treated with two nSMase inhibitors (Inh1 and Inh2) to prevent the formation of TG-containing GMPs. (**F, G**) *In vitro* (**F**) and *in vivo* (**G**) effects of nSMase inhibitors (**F, G**: Inh1 or Inh 2), or platelet (**G**: ‘-Plt.’), or platelet and granulocyte (**G**: ‘-Plt./Gr.’) depletions on the ability of GMP-containing supernatant (**F**) or AMS (**G**) to cause upregulation of *Arg1* in macrophage cell line. For (**D-G**): **, p<0.01; data represented as mean ± s.e.m.; see **Table S1** for statistics. <u>Abbreviations</u>: Gr., granulocytes; GMPs, granulocyte-derived microparticles; nSMase, neutral sphingomyelinase; Plt., platelets; TB, thrombin; TG, triglycerides; Untr., untreated. See also **Table S1** and **Figure S5, Supplemental Material**.

### GMPs contain TGs and FAs

The findings from the above-described experiments suggest that during blood coagulation, murine granulocytes release lipids that are subsequently loaded onto lipoproteins such as HDLs. Platelets are important for activating granulocytes during blood coagulation, ^18^ but we found that GMPs were almost tenfold more potent than platelet-derived products to induce *Arg1* expression (**Figure 4F**). To identify the most active Arg1-inducing lipids, we performed large-scale untargeted lipodomics (829 tested lipids) to compare the lipid compositions of GMPs, AMS, and CMP (622 detected lipids and 38 subclasses, **Figure S6A**, **Supplemental Material**) and found that there were significant differences between the lipid compositions of GMPs vs. CMP and those of AMS vs. CMP (**Figure 6A, B**, and **Figure S6B**, **Supplemental Material**). Compared with CMP, GMPs contained higher concentrations of various TGs consisting of saturated (C15:0, C16:0, C18:0) and unsaturated (C18:1n9c, C18:2n3, C18:2n6, C20:3n3 and C20:4n6) FAs and their metabolic products, such as diglycerides, free (mostly branched) FAs, and carnitine (**Figure 6A**). Compared to CMP, AMS contained high concentrations of lysophospholipids (Lyso-PLs), namely LPA, LPI, LPE, and LPC, and also free FAs (**Figure 6B**). We performed additional targeted analysis of free FAs and found that GMPs contained saturated C14:0, C16:0, and C18:0 FAs (**Figure 6C**), whereas AMS had higher concentrations of saturated C15:0, C16:0, and C18:0 FAs (**Figure 6D**) and unsaturated C18:1n9c, C18:2n3, C18:2n6, C20:3n3, and C20:4n6 FAs than CMP (**Figure 6E**). Overall, compared with CMP, AMS had 30%–90% higher concentrations of saturated free FAs (except for C14:0 FA), 40%–90% higher concentrations of unsaturated free FAs, and 70% higher total concentrations of free FAs (**Figures 6C-E**). Thus, our data demonstrated that there were higher concentrations of free FAs and Lyso-PLs in serum than in plasma and GMPs contained high levels of TGs and free FAs.

**Figure 6.**
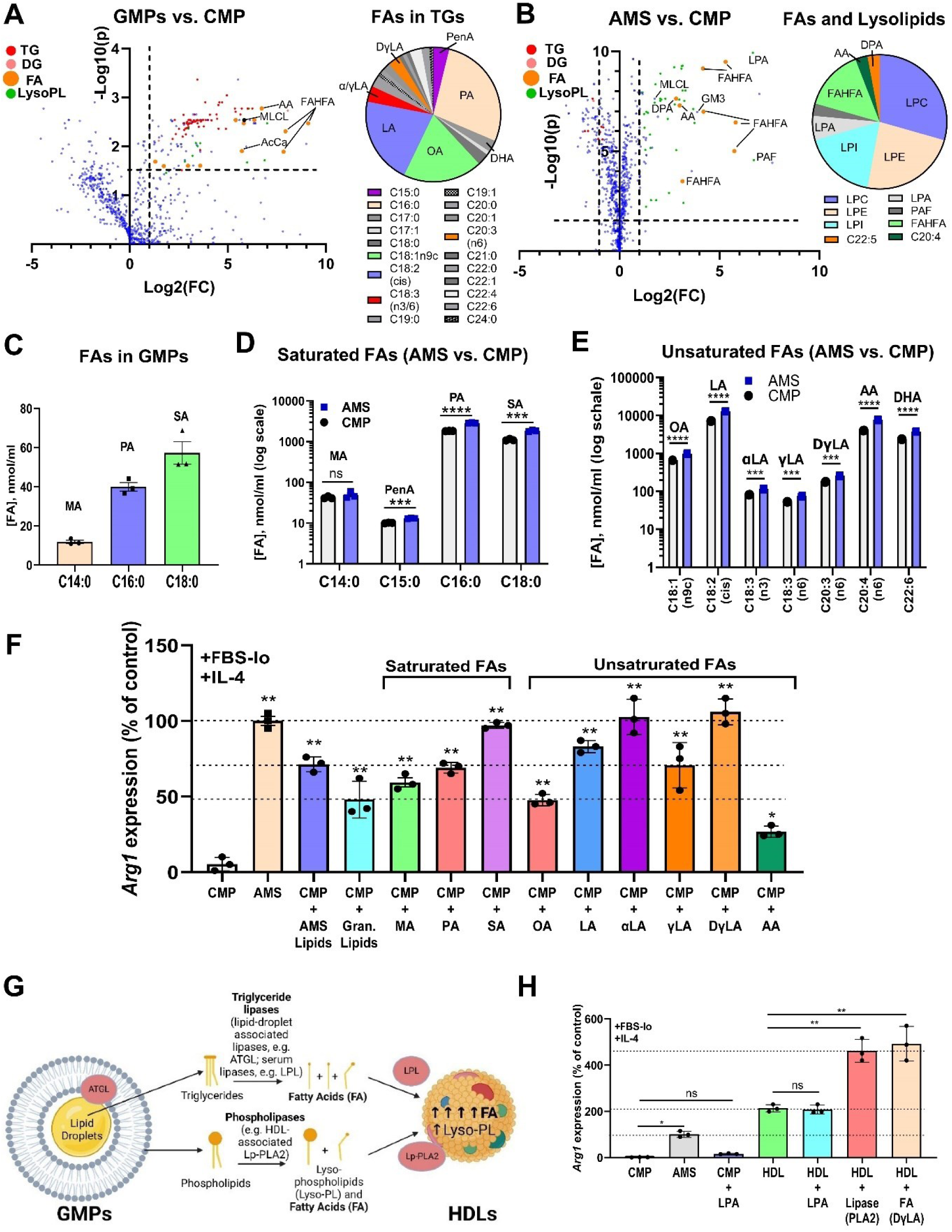
Identification of specific lipids in granulocyte-derived microparticles and adult mouse serum that potentiate M2 activation. (**A**) Global untargeted lipidomics analysis of lipids in granulocyte-derived microparticles compared to citrate mouse plasma (GMPs vs. CMP). The volcano plot is shown on the left panel (x-axis: -log(P-value); y-axis: log(fold change)), while the distribution of fatty acids in GMP-specific triglycerides (marked in red in the upper right quadrant of the volcano plot) is shown on the right panel. (**B**) Global untargeted lipidomics analysis of lipids in adult mouse serum compared to citrate mouse plasma (AMP vs. CMP). The volcano plot is shown in the left panel (x-axis: -log(P-value); y-axis: log(fold change)), while the distribution (left panel) and composition (right panel) of lipids enriched in AMS vs. CMP (FAs, marked in orange color; lysolipids, marked in green color) are shown in the upper right quadrant of volcano plot, and the pie chart, respectively. (**C-E**) Targeted metabolomics analysis of free FAs in GMPs (**C**), and saturated (**D**) vs. unsaturated (**E**) FAs in AMS (blue color) vs. CMP (grey color) indicates a 1.3-2-fold increase in free FA concentrations and substantial decrease in TG concentration in AMS compared to CMP. (**F**) **‘**Gain-of-function’ analysis of AMS- and GMP-derived lipids and various free FAs for their ability to induce *Arg1* in the macrophages. (**G, H**) The role of external lipases on functional activity of HDLs to induce *Arg1* expression in macrophages. (**G**) Scheme of the proposed model of external lipolysis of GMP-derived triglycerides and phospholipids with triglyceride- and phospholipases that are present in granulocyte lipid droplets (e.g. ATGL), serum (e.g. LPL), or associated with lipoproteins such as HDLs (e.g. Lp-PLA_2_). (**H**) Effects of external lysolipids with PLA_2_ or the addition of external fatty acids to HDLs on the ability of these HDLs to induce Arg1 expression in the macrophage cell line. For (**C-F**), (**H**): *, p<0.05; **, p<0.01; ***, p<0.001; ****, p<0.0001; data represented as mean ± s.e.m.; see **Table S1** for detailed statistics. <u>Abbreviations</u>: AA, arachidonic acid; ATGL, adipose triglyceride lipase; FA, fatty acids; Gran., granulocytes; GMPs, granulocyte-derived microparticles; LA, linoleic acid; αLA, α-linolenic acid (ω3); γLA, γ-linolenic acid (ω6); DγLA, Dihomo-γ-linolenic acid; LPL, lipoprotein lipase; Lp-PLA2, lipoprotein-associated phospholipase A2; Lyso-PL, lysophospholipids; MA, myristic acid; OA, oleic acid; PA, palmitic acid; PenA, pentadecanoic acid; SA, stearic acid. See also **Table S1** and **Figure S6, Supplemental Material**.

### Extrinsic FAs induce M2 activation

To perform “gain-of-function” experiments, we tested various lipid compounds found in GMPs and AMS to compare their ability to induce *Arg1* expression with that of CMP. We performed this test by adding lipids extracted from AMS or granulocytes, or purified/synthetic lipids, to CMP (or CBP with comparable efficiency, **Figure S6C**, **Supplemental Material**), which served as a source of murine or bovine lipoproteins. Lipids extracted from AMS and granulocytes exhibited ∼70% and ∼50% of the ability of an AMS-positive control, respectively, to induce *Arg1* expression, whereas pure C18:0 and C18:2n3, C18:2n6, and C20:3n3 demonstrated 80%–100% and C20:4 n6 (arachidonic acid) only 25%–30% of the abilities of the same control to induce *Arg1* expression (**Figure 6F**). Among all other tested compounds, C8:0 octanoic acid exhibited the highest activity, i.e., 110%–120% of the ability of AMS to induce *Arg1* expression. We did not detect C8:0 octanoic acid in GMPs or sera, but this FA is abundant in food supplements such as coconut oil. The other tested lipid compounds, which included multiple lysolipids, TGs (C18:1n9c, C18:1n9c, C18:1n9c), acylcarnitine (C14:0), serum-abundant ganglioside GD3 (**Figure S6D**), and several branched FAs detected in GMPs and AMS (**Figure S6E**), did not exhibit statistically significant abilities to induce *Arg1* expression. We hypothesized that during blood coagulation, TGs and phospholipids from GMPs are degraded by serum- or granulocyte-derived lipases and phospholipases into lysolipids and FAs, respectively (**Figure 6G**). This hypothesis was supported by our finding that treatment of purified and washed HDLs with phospholipase A_2_ (PLA2) – which is normally associated with lipoproteins in sera (Lp-PLA2) – increased their ability to induce *Arg1* expression by a twofold relative to the ability of externally added FA (DγLA) with the high level of activity (**Figure 6H**). In addition, LPLs did not demonstrate any ability to induce the expression of *Arg1* when added to purified HDLs (**Figure 6H**), indicating that FAs but not lysolipids are the components of HDLs that can effectively induce Arginase 1 expression in macrophages. Taken together, our findings show that C18:0, C18:2n3/6, and C20:3n3 FAs mediate the strongest levels of M2 activation during blood coagulation.

### HDL/LDL receptors mediate M2 activation

It was previously established that SR-B1 is an HDL receptor ^27^ and SR-B3/CD36 is an LDL receptor, ^9^ but more recent detailed analyses have demonstrated that both SR-B1 and SR-B3/CD36 can transport FAs with comparable efficiency and thus do not completely discriminate between HDLs and LDLs. ^28,29^ M2 macrophages upregulate the expression of SR-B1 and SR-B3 but not that of other lipoprotein receptors. ^9^ Thus, we blocked SR-B1 or SR-B3 with antibodies (Blocking antibodies, **Figure 7A**) or knocked down SR-B1 or SR-B3 expression with small interfering RNA (siRNA, **Figure 7A**) and observed the results. We found that blockade or knockdown of SR-B1 and SR-B3 in macrophages significantly inhibited Arginase 1 expression (**Figure 7A**). Specifically, antibody blockade of SR-B1 was much more effective than blockade of SR-B3 in inhibiting *Arg1* expression (90 ± 1% and 59 ± 2% reductions in *Arg1* expression, respectively, in the case of ABS (**Figure 7A**, left panel), and 71 ± 5% and 39 ± 8% reductions in *Arg1* expression, respectively, in the case of AMS (not shown)) and siRNA knockdown (50 ± 10% and 30 ± 10% reductions in *Arg1* expression, respectively, in the case of AMS (**Figure 7A**, right panel)). This finding is consistent with the fact that SR-B1 promotes cell growth and survival via the PI3K/Akt pathway, ^30^ which we found to be upregulated by the ABS-hi fraction and a pathway involving cancer, according to a KEGG analysis (**Figure 2D**). Overall, our findings indicate that the HDL receptors play a key role in the uptake of lipoprotein-associated FAs that are required to mediate M2 activation.

**Figure 7.**
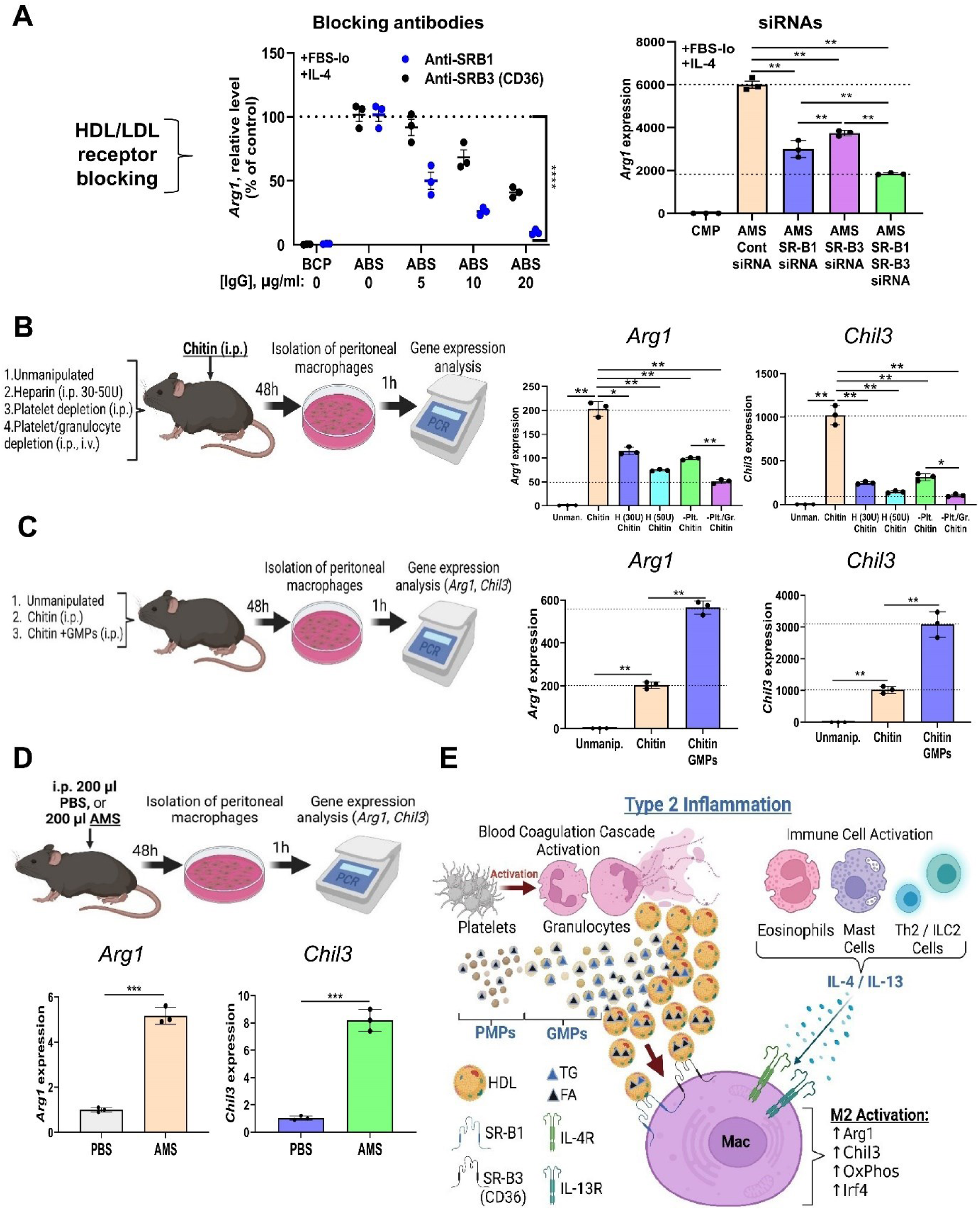
Identification of granulocyte-derived lipids and lipoprotein receptors for IL-4/IL-13 cofactor and their role in type 2 inflammation. (**A**) Analysis of *Arg1* expression in macrophages with inactivated SR-B1 and/or SR-B3 receptors using blocking antibodies (left panel) and siRNA cocktails for receptor knockdown (right panel). (**B**) Experimental design (left panel) and experimental results (right panel) of induction of IL-4/IL-13-mediated type 2 inflammation in peritoneally cavity using i.p chitin administration with or without inhibition of blood coagulation with heparin (H, 30 and 50 U per mouse), or platelet (-Plt.), or platelet and granulocyte (-Plt./Gr.) depletions, and subsequent isolation of peritoneal macrophages for gene expression analysis of *Arg1* and *Chil3*. (**C**) Experimental design (left panel) and observed results (right panel) of induction type 2 inflammation in the peritoneal cavity using chitin with or without the addition of granulocyte-derived microparticles (GMPs), and gene expression analysis of *Arg1* and *Chil3* expression in peritoneal macrophages 48h after co-administration of chitin and GMPs (**D**) Intraperitoneal injection of AMS results in 5-8-fold upregulation of *Arg1* and *Chil3* in peritoneal macrophages. The experimental design is shown in the upper panel, and the observed results are shown on the bottom panel. (**E**) Proposed model of regulation of type 2 inflammation by platelet- and granulocyte-derived FAs associated with microparticles that become loaded on HDLs during blood coagulation when platelets and granulocytes become activated to work synergistically with IL-4 or IL-13 produced by Th2, ILC2, eosinophils, or tissue-resident mast cells. For (**A**)-(**D**): *, p<0.05; **, p<0.01; ***, p<0.001; ****, p<0.0001; data represented as mean ± s.e.m.; see **Table S1** for detailed statistics. <u>Abbreviations</u>: AMS, adult mouse serum; Cont., control; FAs, fatty acids; GMPs, granulocyte-derived microparticles; Gr., granulocytes; H, heparin; PBS, phosphate buffered saline; Plt., platelets; PMPs, platelet-derived microparticles; TG, triglycerides; Unmanop., unmanipulated. See also **Table S1** and **Figure S7, Supplemental Material**.

### The connection between type 2 inflammation and coagulation

Next, we tested whether blood coagulation and granulocytes play a role in vivo during type 2 inflammation induced by intraperitoneal (i.p.) injection of chitin, which is a common model for parasitic or fungal infections and allergies. ^31,32^ First, we observed that mice treated with heparin or depleted of platelets and granulocytes (to prevent blood coagulation processes and the release of granulocyte-derived lipids) and then subjected to i.p. injection of chitin exhibited a substantial decrease in the expression of *Arg1* and *Chil3* in peritoneal macrophages (**Figure 7B**). Second, co-administration of purified GMPs and chitin caused an approximately threefold upregulation in the expression of *Arg1* and *Chil3* in peritoneal macrophages (**Figure 7C**). Third, mice that were not manipulated and were subjected to i.p. injection of AMS exhibited a five-to-eightfold increase in the expression of *Arg1* and *Chil3* in peritoneal macrophages (**Figure 7D**), while the numbers of granulocytes detected in the peritoneal cavity were substantial and comparable to the numbers detected in peritoneal macrophages as early as 24h after chitin administration (**Figure S7**, **Supplemental Material**). These data indicate that HDL–lipid complexes are important cofactors for IL-4 or IL-13 *in vivo* during type 2 inflammation when platelets and granulocytes become activated during blood coagulation that accompanies inflammation and release lipid-containing microparticles. Platelets play an important role in this process in vivo by activating granulocytes during blood coagulation and contributing to an external pool of FAs associated with microparticles (**Figure 7E**).

## Discussion

Our findings demonstrate that external lipolysis during blood coagulation and newly synthesized FAs associated with HDLs play a crucial role in the expression of more than 80% of M2-associated genes, such as *Arg1*, *Chil3*, suppressor of cytokine signaling 2 (*Socs2*), sialic acid-binding immunoglobulin-type lectin F (*Siglec-F*), prostaglandin-endoperoxide synthase 1 (*Ptgs1*), and interferon regulatory factor 4 (*Irf4I*), as well as OXPHOS (**Figure 7E**). Interestingly, we found that the expression of a minority (6 of 32) of very important M2 genes, namely *Mrc1* (which encodes cluster of differentiation (CD) 206), *Mgl2* (which encodes CD301), *Retnla* (which encodes resistin-like molecule alpha), *Egr2* (which encodes an immediate early (IE) gene), *Ccl22* (which encodes C-C motif chemokine 22), and *Klf4* (which encodes the transcription factor Krüppel-like factor 4 downstream of signal transducer and activator of transcription 6), did not depend on lipoprotein-associated FAs in the ABS-hi fraction (**Figure 2A**). It has been found that in mouse macrophages, the expression of CD206, CD301, and RELMα (Retnla) depends on internal lysosomal lipolysis of TGs, ^9^ whereas in human macrophages, the expression of *Mrc1* does not require FA oxidation. ^13^ However, it is technically challenging to block internal sources of FAs and their intracellular oxidation, and it, therefore, remains unclear whether OXPHOS and FA oxidation is required for M2 activation. ^12,33^ Our results indicated the importance of external sources of FAs for M2 activation; these sources are technically easier to block than internal sources and could be used to prove the critical role of FAs in this process. As the ABS-hi serum fraction contained both HDLs and LDLs, lipoprotein- and FA-independent markers were upregulated in our model in the absence of HDLs or LDLs. We hypothesize that this was due to the internal lipolysis of TGs stored in lipid droplets in unstimulated macrophages, ^34^ such that these stores were exhausted rapidly in M2- (but not M1-) stimulated cells. ^9^

We infer from our study that first, the LDL-TG-CD36 pathway ^9^ is most likely important for TG deposition in unstimulated macrophages ^34^ and for the expression of early M2 markers such as *Egr2* (an immediate early gene whose expression is upregulated as early as 3 h post-IL-4 treatment); ^35^ and second, the expression of late M2 markers such as Arginase 1 (whose mRNA and protein concentrations peak on days 4 and 8, respectively; **Figure 1B**) or CCAAT/enhancer-binding protein beta (CEBPβ; a transcription factor downstream of Egr2 ^35^) depends on external lipolysis and HDL–FA complexes, as internal stores of TGs are exhausted as early as 24 h post-stimulation. ^9^ It was shown recently that the expression of RELMα in IL-4-treated murine BMDMs depends on the activity of acetyl-CoA carboxylase, which is responsible for the first step of FA biosynthesis within the cell. ^36^ Future experiments are needed to fully understand the molecular mechanisms of expression of HDL–FA-dependent vs. independent M2 genes.

We believe that the HDL-FA pathway is also very important for the expression of M2 markers and OXPHOS in tissue-resident macrophages in vivo. It was recently reported that lung-tissue macrophages require OXPHOS accompanied by FA oxidation and HDL-mediated cholesterol efflux. ^37,38^ In contrast to LDLs, which are mostly produced in the liver, HDLs can be also produced in tissues such as resting and activated astrocytes in the central nervous system, in which HDL complexes with microRNAs ^39^ and lipids such as FAs; ^40^ these complexes may modulate microglial functions that result in the expression of ARG1 in vivo under homeostatic conditions. ^41^ Microglia also express lipoprotein-FA-dependent CX3C motif chemokine receptor 1 and interferon regulatory factor 4 but not lipoprotein-FA-independent CD206. ^41^ Moreover, alveolar macrophages express Siglec-F, ^38,42^ which requires both lipoprotein–FAs in our model (**Figure 2A**). Further investigations are required to fully characterize the importance of the HDL–FA-dependent pathway for generating the homeostatic phenotype of tissue-resident macrophages.

An interesting finding of this study is that granulocytes (mostly neutrophils) produce large amounts of lipids in GMPs that are subsequently degraded into FAs when loaded onto HDLs or LDLs (in the case of bovine sera). The mechanism of lipid exchange from GMPs to HDL or LDLs has yet to be determined, but it was previously reported that HDLs can form aggregates with microparticles. ^43^ It was also previously demonstrated that platelet-derived microparticles can bind to LDLs, ^44^ and it was later shown that human plasma microparticles can form aggregates with both LDLs and HDLs. ^45^ During this aggregation, lipids from platelet-derived microparticles or GMPs can be loaded onto HDLs and undergo enzymatic conversion from TGs and phospholipids into FAs and Lyso-PLs (**Figure 6G, H**). TG lipases are present in serum ^46^ and lipid droplets of granulocytes, ^47^ whereas PLA2 is associated with lipoproteins. ^48^ HDLs have been shown to play an anti-inflammatory role in macrophages,^39,43,49,50^ but the mechanisms of this process are not clear, and the results are debated. The findings of the current study indicate the importance of HDL-associated lipids, namely the need for specific FAs for anti-inflammatory processes in macrophages, which has not been examined in previous studies.

The blood coagulation cascade, platelets, and neutrophils are crucial for inflammation and tissue repair. ^51–53^ Neutrophils are known to mediate the M2 phenotype of macrophages during parasitic infection and initiate anti-inflammatory pathways via secretory factors, but these factors are unknown. ^54,55^ Moreover, neutrophils arrive at the site of inflammation as early as two hours after injury and remain in tissues for up to five days, ^52^ providing a solid source of FAs for the M1-to-M2 transition by inhibiting the M1 pathway and promoting the M2 pathway. The slow kinetics of Arginase 1 expression especially on a protein level, as shown by the fact that its concentration peaked on day 8 (**Figure 1B**), could also indicate that during acute inflammation, neutrophils initiate *Arg1* mRNA expression as early as 24–48 h after injury, resulting in subsequent increases in ARG1 on protein level and a switch from glycolysis to OXPHOS. Thus, the M1-to-M2 transition is typically completed 5–7 days after injury during acute inflammation. ^52,53^ Neutrophils are also the last population that appears among circulating fetal blood cells. ^56^ This explains why ABS was more potent than FBS in our experiments. Differences between the activities of various batches of FBS may be attributable to the contamination of FBS with ABS and to differences between the stages of development in which the batches of FBS were collected. Overall, our newly discovered granulocyte-mediated pathway indicates that the process of alternative (M2) macrophage activation is highly complex as shown in the scheme in **Figure 7E**, which supports the new concept of a continuous spectrum of activation rather than a simple M1–M2 dichotomy.

## Supporting information

Supplemental Material single PDF file

## Acknowledgments

We thank the Laboratory Animal Service Centre Facility at the Chinese University of Hong Kong (Hong Kong) for assisting with animal maintenance. We also thank the Centre for Panor-Omics Sciences, Lee-Ka Shing Faculty of Medicine, Hong Kong University (Hong Kong) for protein LC-MS and fatty acid GC-MS analyses and the Hong Kong office of BGI company for global untargeted LC-MS lipidomics. English editing of the manuscript was performed using a service provided by AsiaEdit Inc (Hong Kong). Schematic images were made using BioRender software. The work was supported by the Research Grant Council – Areas of Excellence Fund grant, reference no. AoE/M-604/16 (Hong Kong Government, Hong Kong), Faculty Development Competitive Research Grant Program, reference no. 201223FD8829 from Nazarbayev University (Nazarbayev Fund, Kazakhstan), and a Social Policy Research Grant from Nazarbayev University (Nazarbayev Fund, Kazakhstan).

## Author contributions

Conceptualization: T.V. and E.D.P.

Methodology: T.V. and E.D.P.

Investigation: T.V. and E.D.P.

Data analysis: E.D.P. and T.V.

Funding acquisition: E.D.P.

Project administration: T.V. and E.D.P.

Supervision: E.D.P.

Chemical synthesis and purification of L-13-DHAHLA and R-13-DHAHLA lipids: A.V. and T.V.H.

Writing – original draft: E.D.P. and T.V.

Writing – review & editing: E.D.P., T.V., A.V. and T.V.H.

## METHODS

### Mice

C57BL/6 (B6), B6.129S4-C3tm1Crr/J (C3^-/-^), and B6.129S4-C4btm1Crr/J (C4^-/-^) mice were purchased from Jackson Laboratories and maintained at the Laboratory Animal Service Centre, the Chinese University of Hong Kong. Experimental procedures were approved by the Health and Food Department of the government of Hong Kong, and Animal Experimentation Ethics Committee at the Chinese University of Hong Kong, and the City University of Hong Kong. Genotyping and phenotyping of C3^-/-^ and C4^-/-^ animals were performed by standard PCR and western blotting, respectively as described earlier in our previous studies. ^57^ The study was performed following the recommendations of the ARRIVE guidelines. Animals were housed under a controlled environment to maintain constant temperature and humidity with an automatic 12-hour light and dark cycle. Animals were provided with housing cages with 4–5 animals of the same sex per cage and were fed ad libitum. The bedding was replaced every 2-3 days and materials including bedding were passed through decontamination and/or sterilization procedures before entering the animal facility. From 6 to 12-week-old male and female animals were used in our study for bone marrow isolation (6-8 weeks old) and other experiments (8-12 weeks old). Bone marrow was isolated by mixing equal numbers of femur bones from male and female animals.

### Cell lines

Mouse macrophage RAW264.7 (TIB-71™) cell line was purchased from the American Tissue Culture Collection. The cells were used for no more than 10-15 passages, and authentication was performed by morphology and flow cytometry for the expression of macrophage markers CD11b and CD86 as was reported earlier. ^58^ Cells were grown in DMEM (Gibco 11965118) supplemented with 10% fetal bovine serum (FBS, Gibco 10082-147) and 100 U/ml penicillin– streptomycin (Gibco 15140122) as was described in our previous studies. ^35^ The cells were regularly tested by PCR and were negative for Mycoplasma contamination. For experiments, adherent cells incubated in DMEM with 10% FBS were washed three times in serum-free DMEM media and incubated in serum-free condition or supplemented with a 10% ‘FBS-low’ (less than 100 kDa) fraction of FBS.

### Cell isolation and culture

Bone marrow–derived macrophages (BMDM) were grown in DMEM supplemented with 10% FBS and 100 U/ml penicillin–streptomycin in the presence of macrophage colony-stimulating factor (M-CSF, 10 ng/ml, R&D Systems 416-ML-050), as described earlier in our studies. ^39,59^ Bone marrow from 6- to 8-week-old B6 mice were isolated by flushing with PBS from femur bones, and single-cell suspensions were obtained by passing bone marrow through a 70-μm cell strainer (Falcon). For experiments, BMDM were grown in DMEM with 10% FBS and 100 U/ml penicillin–streptomycin for five days, adherent cells were washed three times in DMEM and then incubated in serum-free DMEM, or DMEM supplemented with 10% ‘FBS-lo’ fraction of FBS. Flow cytometry analysis (see below) indicated 97–99% purity of the population of CD11b^+^F4/80^+^ cells. ^59^ Peritoneal macrophages (PMs) were isolated from 8–12-week-old B6 mice by peritoneal lavage with 10 ml of PBS, centrifugated at 300g for 10 min, resuspended in DMEM with 10% FBS and 100 U/ml penicillin–streptomycin, and allowed to adhere to plastic for 8h at CO_2_ incubator similarly as we did earlier. ^59^ After adherence, non-adherent cells were removed by washing plates with PBS without calcium and magnesium leading to a 95-98% pure population of adherent cells (PMs) as determined by flow cytometry for expression of CD11b and F4/80. ^59^

### Specific reagents

The following lipid-related reagents were used in our study: 1) *fatty acids*: docosapentaenoic acid (DPA n3, 22:5 n3, Cayman 90165), osbond acid (DPA n6, C22:5 n6, Cayman 10008335), docosahexaenoic acid (DHA, C22:6 n3, Cayman 90310), arachidonic acid (AA, C20:4 n6, Cayman 90010), stearic acid (SA, C18:0, Cayman 10011298); oleic acid (OA, C18:1n9c Cayman 90260), linoleic acid (LA, 18:2 n6, Cayman 90150), α-linolenic acid (αLA, C18:3 n3, Cayman 90210), γ-linolenic acid (γLA, C18:3 n6 Cayman 90220), dihomo-γ-linolenic acid (DγLA, C20:3 n6, Cayman 90230), palmitic acid (PA, C16:0, Cayman 10006627), myristic acid (MA, C14:0, Cayman 13351), octanoic acid (OctA, C8:0, Cayman 33674); 2) *branched fatty acid esters of hydroxy fatty acids* (FAHFAs): 13-POHSA (Cayman 17111), 9-PAHSA (Cayman 17037) 13L-DHAHLA and 13R-DHAHLA were synthesized as described earlier ^60^; 3) *lysolipids*: lysophosphatidic acid sodium salt (LPA, Abcam ab146430), lysophosphatidylcholine (LPC, Cayman 24331), lysophosphatidylethanolamine (LPE, Cayman 25844), lysophosphatidylinositol (LPI, Cayman 26016), platelet activating factor (PAF, Cayman, 60900); 4) *triglicerides*: oleic acid triglyceride (TG) was purchased from Sigma (T7140); 5) *carnitines*: myristoil-L-carnitine (Cayman 26559) 6) *lipases*: phospholipase A2 (PLA2) from honeybees was purchased from Sigma (P9279). For functional assays, lipid reagents were mixed with CMP, or CBP, or purified HDLs before being added to cultures.

### Serum and plasma preparations

Heat-inactivated fetal and adult bovine serums (FBS and ABS, respectively) were purchased from Gibco. Adult bovine plasma with sodium citrate and adult bovine serum were purchased from Innovative Research. Normal (NHS) or C3- (A314), C4- (A308), and C3/C4- (A340) depleted human adult serums were purchased from Complement Technology Inc. To obtain adult mouse serum (AMS), blood was collected from B6 (wild-type, WT), C3^-/-^ or C4^-/-^ anesthetized animals via cardiac punctures and allowed to coagulate at 37 C for 1h. The blood clot was removed after low-speed centrifugation (300 g, 10 min) and the serum was heat-inactivated (56 C, 30 min). Mouse citrate plasma (MCP) was obtained from B6 mice using 0.38% sodium citrate (Sigma) as an anticoagulant. The serums were centrifugated at high speed (20,800 g), passed through a 0.2µ filter to remove debris, and separated into two ‘hi’ (more than 100 kDa) and ‘lo’ (less than 100 kDa) fractions using 100 kDa filters for 50 ml and 1.7 ml tubes (Vivaspin 20 and 500, Sartorius). Bovine serum albumin (BSA, fraction V) and bovine immunoglobulins (IgG fraction) were purchased from Sigma.

### M1 and M2 activation

M1 and M2 activations of BMDM or RAW264.7 cells were performed in serum-free DMEM with 100 U/ml penicillin–streptomycin, or DMEM supplemented with 10% ‘FBS-lo’ fraction in the presence of other serum fractions (e.g. ‘ABS-hi’) at the quantities of 10% (v/v). For M1 activation, the cells were incubated with IFNγ (100 ng/ml, R&D Systems 485-MI-100) and LPS (100 ng/ml, Sigma L5293) for 24h as done earlier. ^35^ For M2 activation, the cells were incubated with IL-4 (50 ng/ml, Peprotech 214-14) or IL-13 (50 ng/ml, Peprotech 210-13) for 24h or 4 ^35^, 8, or 10 days.

### RNA isolation and real-time RT PCR

For RNA isolation, the adherent RAW264.7 cells, BMDMs, or PMs were washed with PBS and then lysed in a culture plate using QIAzol Lysis Reagent (Qiagen 79306) as was described in our previous studies. ^35,57,59,61^ RNA purification was performed using a DNase digestion kit (RNase-Free DNase Set, Qiagen 79254), miRNeasy Mini kit for RNA isolation (Qiagen 217004), and nuclease-free water (Life Technologies AM9937) for RNA dilution. cDNA was synthesized using TaqMan™ Reverse Transcription Reagents (Life Technologies N8080234). Real-time RT PCR was performed using ABI ViiA 7 and ABI QuantStudio 7 Flex Systems with MicroAmp® Optical 384-Well Reaction plates with specific barcodes (Life Technologies 4309849) and Power SYBR™ Green PCR Master Mix (Life Technologies 4368708). The primer sets are summarized in the **Table. S2**, **Supplementary Material**. Relative expression levels were calculated using the ΔΔC_T_ method and normalized to the expression of the GADPH housekeeping gene, and then to the expression of the control sample, which was defined as 1 or 100%.

### Protein electrophoresis and western blotting

The cells were lysed in RIPA buffer with Halt™ Protease Inhibitor cocktail and then diluted in Bolt™ LDS sample buffer (all reagents are from Thermo Fisher Scientific) with or without 10 mM dithiothreitol (DTT, Sigma) treatment for the reduction of cysteines. The samples were run on Bolt^TM^ bis-tris plus 4-12 % gels using Bolt™ MES SDS running buffer (Thermo Fisher Scientific NW04122Box and B0002, respectively) and Precision Plus Protein™ dual color standards (Bio-Rad 1610394) at a voltage of 200V for 25 minutes. The gel staining for protein content was done using Pierce™ silver stain/distain kit (Thermo Fisher Scientific 24612). Immunoblot analysis was performed using PVDF membranes (Pierce 88518) and blotting-grade blocker (Bio-Rad 1706404) of nonspecific absorbance according to standard protocols as previously reported in our earlier studies ^35,39,61^. The antibodies to the following antigens were used for western blot analysis: β-Actin (Cell Signaling 4967), ARG1 (Cell Signaling 9819), CHIL3 (Biolegend 853803), NOS2 (Cell Signaling 2982), and APOA1 (Biomatik CAC08238). As a secondary antibody, we used anti-rabbit antibodies conjugated with horse radish peroxidase (HRP, Cell Signaling 7074S). As substrate for HRP, we used an ECL high-sensitivity substrate kit (Abcam ab133406) and Kodak film. For protein level assessments in lysed cells, β-Actin was used as a loading control, and quantitative analysis was performed using ImageJ software v. 1.8.0. For protein concentration assessment of fractionated serum fractions, we used Bradford assay (Sigma).

### Arginase enzymatic activity assay

Arginase enzymatic activity was measured in BMDM cell lysate using a colorimetric Arginase Activity Assay kit (Abcam ab180877) according to manufacturers’ protocol.

### RNA-seq and data analysis

Whole-transcriptome sequencing analysis of untreated BMDMs, or treated with IL-4 and/or ABS-hi serum fraction was performed at the Centre for Panoromics Sciences, Lee-Ka Shing Faculty of Medicine, Hong Kong University (Hong Kong) similarly as we published earlier. ^62^ RNA samples from the cells in three separate wells were isolated as described above (see RNA isolation section) and then passed through the quality control procedure that included concentration and purity assessment by UV spectrophotometry (260/280 nm absorbance ratio), fluorometric quantification using RNA-specific fluorescent probe (Qubit Flex Fluorometer, Thermo Scientific), and Bioanalyzer RNA quality assessment. cDNA libraries were then prepared from RNA samples using the KAPA-stranded mRNA-Seq kit (Roche KK8421). The sequencing was performed using the Illumina NovaSeq 6000 system. The raw sequence data were further filtered to remove ribosomal RNA and low-quality reads and then reads of more than 40bp were analyzed to align with the mouse genome database. Differential gene expression (DEG) analysis and signaling pathway analysis of four groups of untreated BMDM incubated in DMEM with 10% FBS-lo or treated with IL-4, ABS-hi, or IL-4 and ABS-hi was performed using the DESeq2 R software package (v. 1.26.0) and KEGG software.

### Serum fractionation and characterization

Adult bovine serum (ABS) was diluted in PBS (Gibco), centrifugated at high speed (20,800 g), passed through a 0.2µ filter to remove microparticles and debris, and ABS-hi fraction (more than 100 kDa) was separated by size exclusion as described above (see Serum and Plasma Preparation section). The ABS-hi fraction was further analyzed by gel filtration using Sephacryl S-200 HR and C16/40 Columns (both from Cytiva). Each 1 ml collected fraction was analyzed for protein concentration using a Nanodrop spectrophotometer (absorbance at 280 nm) and pooled three fractions were further analyzed for protein concentration using Bradford assay (Sigma) and used to measure biological activity to induce *Arg1* expression in RAW264.7 macrophage cell line. The active fractions after gel filtration were analyzed for bovine APOA1 and APOB content by ELISA kits (LS Bio LS-F25212 and Novus biologicals NBP2-77030, respectively), and were found positive for APOA1 and negative for APOB (not shown). The biologically active fractions were then further analyzed on Pierce™ Strong Anion Exchange Spin Columns (Thermo Scientific 90011) according to the manufacturer’s protocol. After anion exchange columns, the fractions were eluted with 0.25 M NaCl and biologically active fractions were further analyzed on a hydrophobic interaction column (Macro-Prep® t-Butyl HIC Resin, Bio-Rad 1580090). Three fractions eluted with buffers with pH=7, pH=9, and 8M Urea were further analyzed for their biological activity and APOA1 content by western blotting.

### Protein mass spectrometry

After the electrophoresis and silver staining procedures described above (see Protein Electrophoresis section), the gels were rinsed several times with water, and two bands (Fig. S5a, band 1 and band 2) were dissected. The sliced gel samples were distained using Pierce™ silver stain/disdain kit (Thermo Fisher Scientific), and in-gel digested with trypsin (Sigma). Protein digestion fragments (peptides) were then extracted from the gel and analyzed on Thermo Fisher Scientific LTQ Velos LC-Mass Spectrometer at Bioscience Central Research Facility, Hong Kong University of Science and Technology (Hong Kong). The obtained m/z peaks were identified on the MASCOT server.

### Real-time cell metabolic assay

Oxygen consumption rate and media acidification were measured in unstimulated BMDM incubated in DMEM supplemented with 10% FBS-lo fraction or M2-stimulated cells treated with IL-4, ABS-hi, or IL-4 and ABS-hi using Extracellular Flux Assay kits (XFE96 Flupax mini 102601-100 kit, Seahorse XF DMEM assay medium pack 103680-100 kit, XF cell mitostress test 103015-100 kit; all from Agilent Technologies) and Seahorse XF Pro Analyzer according to manufacturer’s recommendations and previously published protocol. ^63^

### Lipoprotein isolation

Bovine and mouse VLDL/LDL, HDL, and other (non-lipoprotein) ‘protein’ fractions were isolated by NaBr density gradient ultracentrifugation using CS-150GXII Micro-Ultracentrifuge (Hitachi) with S140-AT Rotor (Thermo Scientific) according to the available protocol for S140-AT rotor with modifications. ^64^ Since mouse and bovine serums contained very low levels of VLDLs (not shown), LDL and VLDL fractions were not separated. VLDL/LDL (further referred to as ‘LDL’) fraction was separated from HDL proteins by mixing 600 µl of ABS or AMS with 300 µl of 0.195 mol NaCl and 2.44 mol NaBr (1.006<ρ<1.0063 g/m) and centrifugation at 1,048,600 g, 16 C, 80 min. After removing 300 µl of upper LDL fraction, we added 300 µl of 0.195 mol NaCl and 7.65 mol NaBr (1.0063<ρ<1.21 g/m) and centrifugate at 1,048,600g, 16 C, 140 min. After centrifugation upper 300 µl of HDL fraction was collected, 300 µl of the middle fraction was then discarded, and the bottom 300 µl fraction was collected as a ‘protein’ fraction. HDL, ‘LDL’, and ‘protein’ fractions were then washed three times with PBS using a 100 kDa filter (Vivasipn).

### Enzymatic treatments of HDLs

Before isolation of HDLs, AMS was left untreated or treated with Proteinase K (50 U/ml), trypsin (0.25%), β-Galactosidase from *Saccharomyces fragilis* (40 U/ml), and Neuraminidase (15 U/ml) (all from Sigma) for 2h at 37 C similarly as we did earlier for lipid raft particles. ^65^ Isolated HDLs were then washed with PBS using a 100 kDa filter before adding to the macrophage cell line. For phospholipase A2 (PLA2, Sigma P9279) treatment, isolated HDLs were washed with PBS with EDTA (Gibco), treated with PLA2 from honeybees (see Special Reagent section) for 1h at 37C, and then washed three times with PBS using a 100 kDa filter (Vivaspin).

### Lipid isolation and liposome construction

Lipids were isolated from AMS using Folch lipid extraction protocol^66^. For liposome construction, extracted lipids were dried overnight using a SpeadVac concentrator (Thermo Fisher Scientific), reconstituted in PBS, sonicated for 1h, and vortexed for 20 min as we did earlier. ^65^

### Flow cytometry

The single cell suspensions were prepared from blood, bone marrow, peritoneal cavity, or cultured cells stained for the expression of surface markers with specific antibodies in PBS with 5% FBS and 20% normal goat serum (Jackson ImmunoResearch 005-000-121) and 0.01% NaN_3_ for 20 min on ice. FcRs were blocked using purified anti-mouse CD16/CD32 (2.4G2 101302, 1:50 dilution) antibodies. After staining, the cells were washed with PBS and fixed with 1% paraformaldehyde (Sigma) in PBS. The following antibodies were used CD11b-AF488 (M1/70 101217), Ly6G-PE (1A8 551461), F4/80-APC (BM8 17-4801-82), CD45-APC-Cy7 (30-F11 103116), CD61-FITC (2C9.G2 104306), CD41-PE (MWReg30 133906), TER119-APC (TER-119 116212), CD3ɛ-FITC (145-2C11 100305), CD19-PE (6D5 115502). Antibodies were purchased from Biolegend, BD Biosciences, and eBioscience. The following cell populations were identified according to cell marker expression: macrophages (CD11b^+^F4/80^+^Ly6G^-^), granulocytes (CD11b^+^F4/80^-^Ly6G^+^), mononuclear leukocytes ‘lymphocytes’ (CD45^+^Ly6G^-^), T and B cells (CD3^+^CD45^+^ and CD19^+^CD45^+^, respectively), platelets (CD61^+^CD41^+^), and erythrocytes (TER119^+^). Live/dead cells were discriminated according to FSC/SSC parameters and this gating strategy was confirmed by staining with Live/Dead viability dye e506 (eBioscience). For analysis of microparticles, FSC and SSC parameters were set to a logarithm scale with the decreased threshold for the FSC parameter (1,000) as we did earlier ^53^ Data was collected on BD LSRFortessa^TM^ flow cytometer (BD Biosciences) and analyzed using FlowJo software v. 10.8.0 (Tri Star).

### Isolation of populations of blood cells

Blood was collected from anesthetized B6 mice via cardiac punctures using 0.38% sodium citrate as an anticoagulant. A pooled blood sample (3 ml) was centrifugated at low speed (300 g, 5 min), platelet-rich plasma was collected, and platelets were centrifugated at high speed (20,000 g, 10 min) and resuspended in 3 ml of PBS with calcium and magnesium (initial volume of blood) as we did earlier to obtain washed platelets. ^62^ The cell pellet was then diluted with the same volume of PBS as the volume of collected plasma, overlayed on 5 ml of 70% Percoll (100% Percoll was prepared by mixing 9:1 (v/v) Percoll pH 8.5-9.5 from Sigma with 10X PBS pH=7.4, Gibco), and centrifugated at 500g for 30 min, 20 C, without brake. A population of mononuclear leukocytes (referred to as ‘lymphocytes’ since this population contains >90% of T/B cells as determined by flow cytometry, not shown) was collected on a top of 70% Percoll gradient, washed one time in PBS, and finally resuspended in 3 ml of PBS with calcium and magnesium. The bottom fraction of cells was washed with 3 ml of PBS, overlayed on top of 5 ml of 86% Percoll gradient, and centrifugated at 500g for 30 min, 20 C, without brake. The upper fraction on top of 86% Percoll gradient (granulocytes) was separated from the bottom fraction (erythrocytes), washed, and each fraction was resuspended in 3 ml of PBS with calcium and magnesium. The purities of populations of platelets, mononuclear leukocytes (‘lymphocytes’), granulocytes, and erythrocytes were assessed by flow cytometry and constituted 93-96% (not shown). The cells in each population were stimulated with thrombin (TB, 0.5 U/ml, Life Technologies RP43104) for 30 min at room temperature (20-25 C). After stimulation, granulocytes, lymphocytes, and erythrocytes were centrifugated at a low speed (300 g, 10 min), and supernatant alone, or mixed with CMP or CBP (1:1, v/v), was added to the macrophage cell line for *Arg1* expression analysis.

### Granulocyte isolation and activation

Mouse granulocytes were isolated from bone marrow as in the published protocol ^22^ with minor modifications. Red blood cells were lysed in single-cell suspension of bone marrow cells using ACK Lysing Buffer (Thermo Fisher Scientific) and granulocytes were separated from mononuclear cells using 70%/86% Percoll gradient at interphase. The cells were further purified with magnetic beads and LS Columns from a Mouse neutrophil isolation kit (Miltenyi Biotec 130-097-658) resulting in 97-99% purity as determined by flow cytometry. Granulocytes (10^6^ cells per ml) were left untreated, or pre-treated with neutral sphingomyelinase Inhibitor 1 (3-O-Methyl-sphingomyelin, Abcam ab141756, 50 µM) or Inhibitor 2 (GW 4869, Cayman 13127, 10 µM) for 30 min, washed with PBS without calcium and magnesium, resuspended in the same as the initial volume of PBS with calcium and magnesium (Gibco), and stimulated with thrombin (TB, 0.5 U/ml) for 30 min at room temperature (20-25 C).

### Granulocyte-derived microparticles

Analysis of granulocyte-derived microparticles (GMPs) was performed by flow cytometry and triglyceride release assay was determined in supernatant after low-speed centrifugation (300 g, 10 min) using Triglyceride Assay kit (Abcam ab65336). GMPs were collected in a supernatant of granulocytes (3-5 x 10^6^/ml) treated with thrombin (TB, 0.5 U/ml, 30 min) after low-speed centrifugation. For in vivo experiments, GMPs were washed from thrombin by centrifugation at high speed (24,000g, 15 min) and resuspension in 1 ml of sterile PBS.

### In vivo treatments

Neutral sphingomyelinase Inhibitor 1 (3-O-Methyl-sphingomyelin, 20 µg per mouse) or Inhibitor 2 (GW 4869, 10 µg per mouse) were injected i.v. 25-30 min before blood collection. Platelet and granulocyte depletions were performed as described earlier ^65,67^ resulting in 70-80% of platelet and 50-60% of granulocyte depletion as determined by cell count multiplied by the percentage of cells determined by flow cytometry (not shown).

### Receptor blocking

SR-B1 and SR-B3 receptors were functionally inactivated by using blocking anti-SR-B1 and anti-SR-B3/CD36 antibodies (both used at initial IgG concentration of 40 µg/ml, Thermo Fisher Scientific, PA3-16794 and MA5-16831, respectively) at serial dilutions 1:4, 1:2, and 1:1 (v/v). Alternatively, the knockdown of SR-B1 and/or SR-B3 was performed using a cocktail of four siRNAs. Antibodies were added to the cells 30 min before the addition of ABS-hi fraction and IL-4. Cocktails of four interfering RNAs (siRNAs) for SR-B1 (siGenome smart pool, Mouse Scarb1 siRNA 20778), SR-B3/CD36 siRNA (siGenome smart pool, Mouse CD36 siRNA 12491), and control siRNA (siGenome non-targeting pool, Negative Control siRNA D-00112061305) were purchased from Dharmacon and used similarly as described earlier in our studies. ^35^ BMDMs were transfected with siRNAs using the TransIT-X2 system according to the manufacturer’s protocol. The efficiency of transfection was 60-80% as determined by Cy3-labeled reporter RNA (Life Technologies) in CD11b^+^F4/80^+^ macrophages as determined by flow cytometry (not shown).

### Type 2 inflammation

Chitin from shrimps (Sigma C7170-100G, 1 µg per mouse) was washed three times with sterile PBS, passed several times through 21g needle, and injected i.p. alone or with heparin (Sigma H3149-25KU, 30 or 50 U per mouse), or with washed GMPs (1 ml per mouse, see GMP preparation above) 48h before isolation of peritoneal macrophages and their analysis for the expression of *Arg1* and *Chil3*. AMS was injected i.p. (200 µl per mouse) 48h before isolation of peritoneal macrophages.

### Global untargeted lipidomics

An unbiased global lipidomics analysis was performed by BGI Inc. (Hong Kong Office, Hong Kong). Samples (CMP, AMS, and GMPs; n=5) were frozen and sent to the company on dry ice within 2-4 hours. After thawing at 4 C, 100 µl of supernatant was transferred to a 96-well microplate, and 300 μl of isopropanol (-20 C pre-chilled) was added with internal standards solution (10 µl). The mixture was then vortexed for 1 min and kept at -20 C overnight. The samples were then centrifuged at 4 C for 20 min (800 g) and supernatant was used for LC-MS analysis. To each analyzed sample, 10 μl of quality control reagents were added for quality control assessment and evaluation of the repeatability and stability of analyzed samples. An LC-MS system (Waters 2D UPLC, CSH C18 column (Waters) and high-resolution mass spectrometer Q Exactive (both from Thermo Fisher Scientific)) was used for data acquisition in positive-ion and negative-ion mode respectively to improve the lipid coverage. LC-MS/MS data were analyzed using LipidSearch v. 4.1 software to perform intelligent peak alignment, lipid identification, and calculation of the peak area. LipidSearch database contains information about 1.7 million lipid ions and their predicted fragment ions in eight categories and 300 subclasses, with a total number of 829 lipids identified in our experiments. BGI’s metabolomics software package MetaX was used for further statistical analysis. Principal component (PCA) analysis was used to observe the intergroup differences of all groups of five samples. Differential lipid composition in AMS vs. CMP and GMP vs. CMP was performed by combining with fold change (FC) analysis (log_2_FC is shown), and q-value that was obtained by false discovery rate correction of the p-value from Student’s t-test (-lg[q-value] is shown). The criteria for differential expression of lipid molecules were as follows: FC≥1.2 (up) or FC (down) ≤ 0.83, and q<0.05.

### Targeted metabolomics of free fatty acids

Targeted quantitation of free C4-C26 saturated and unsaturated fatty acids was performed by a combination of gas chromatography-mass spectrometry (GC-MS) at Centre for Panoromic Sciences - Proteomics and Metabolomics Core, Li-Ka Shing Faculty of Medicine, University of Hong Kong (Hong Kong). Supelco 37-component fatty acid methyl ester (FAME) mix, the internal standards (19:0 FAME and all fatty acids) were purchased from Sigma. Samples (CMP, AMS, and GMPs) were frozen and sent to the facility on dry ice within 2 hours after preparation. To make FAMEs, 50 µl of chloroform with 20 µg C19:0 fatty acid internal standard was added to the analyzed samples. The samples were incubated with 0.5M and then 1M HCl at 110°C, and then separated by liquid-liquid extraction using hexane (Sigma). The hexane extract was then dried under a fume of nitrogen at 45 C. For further esterification, 1 ml of methanol and 50 µl of concentrated hydrochloric acid (35%, w/w) were added to the samples under nitrogen fume. After mixing, the tube was incubated at 100 C for 1.5 h and allowed to cool down to achieve 20 C for 30-40 min. After which, 1 ml of hexane and 1 ml of water were added for the final FAME extraction solution. The tube was then vortexed, and after phase separation, 1 µl of the hexane phase was injected for GC-MS analysis. GC-MS chromatogram was acquired using Agilent 7890B GC - Agilent 7010 Triple Quadrupole mass spectrometer (Agilent Technologies Inc). Data acquisition and analysis were performed using the Agilent MassHunter software.

### Statistics

Statistical analyses were performed using GraphPad Prism v. 10.0.2 software. A two-tailed unpaired Student’s t-test was used to determine significant differences between two independent groups. One-way analysis of variance (ANOVA) with Tukey’s post-hoc tests was used to determine significant differences for comparisons among three or more experimental groups with one variable parameter. Two-way ANOVA was used for comparisons between two experimental groups with two variables ([gene expression level] X [time]). P-values of less than 0.05 were considered to be significant with *p < 0.05, **p < 0.01, ***p < 0.001, ****p < 0.0001 shown. Detailed statistical analysis with F, T, and P values is presented in **Table S1**, **Supplemental Material** for main figures and figure legends for supplementary figures.

