## Supplemental Material single PDF file for "Alternative Macrophage Activation Requires Granulocyte-Derived Lipids"

**The PDF file includes:**

**Figures S1-S10 with legends**…pages 2-14

**Tables S1-S2**…pages 15-19

### SUPPLEMENTAL FIGURES

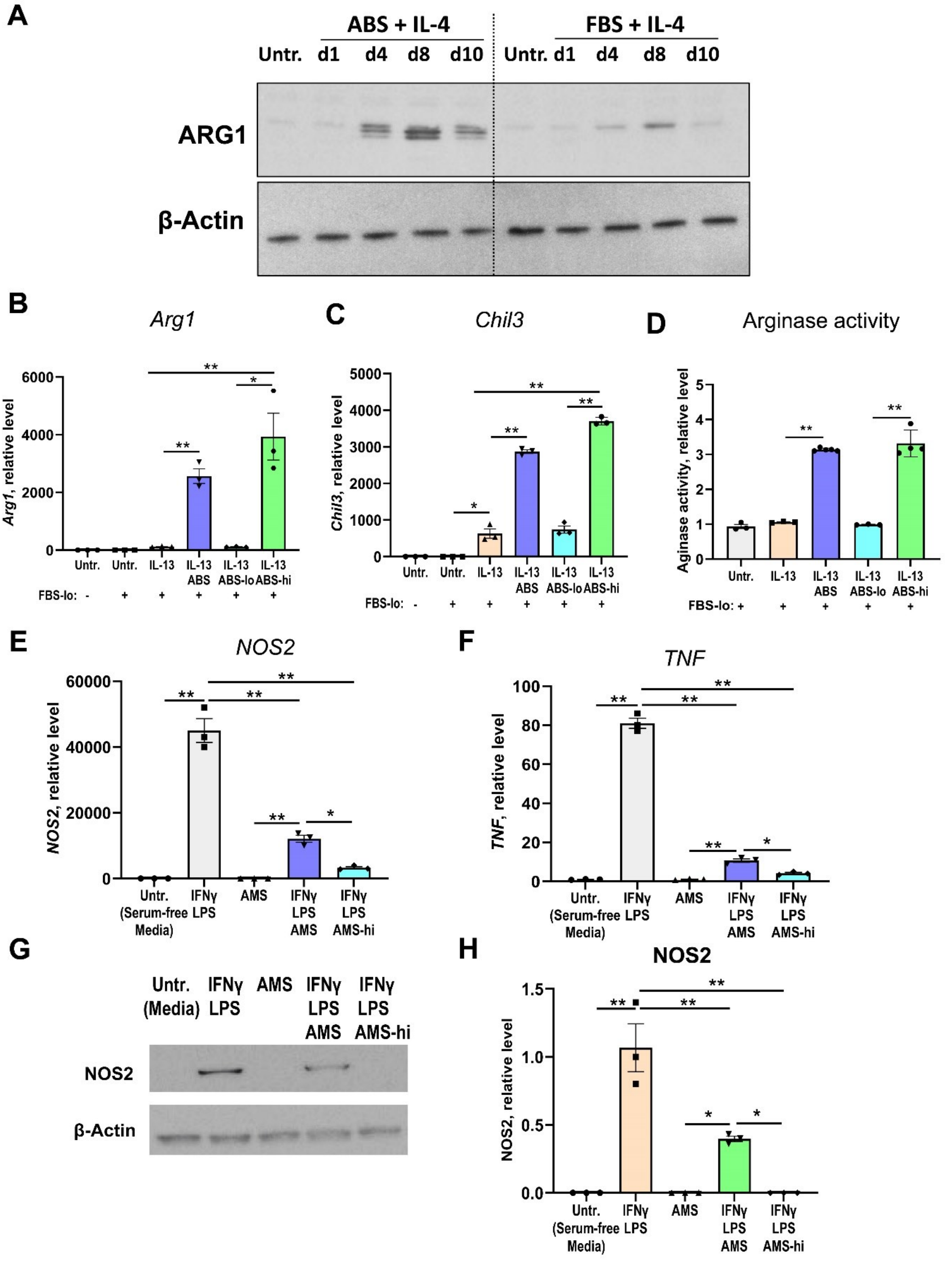


**<u>Figure S1</u>. The influence of serum cofactor on the expression of M2 or M1 markers in IL-4/IL-13- or IFNγ/LPS- treated macrophages.**

(**A**) Effects of FBS and ABS on the kinetics (day 1, 4, 8, and 10) of the expression of Arginase 1 on protein level (ARG1, see Methods) in IL-4 - treated or untreated murine BMDM. A representative western blot image is shown, and the quantification is shown in **Figure 1B**, right panel.

(**B-D**) Comparisons of effects ABS, ABS-lo (<100 kDa), and ABS-hi (>100 kDa) adult bovine serum fractions on the expression of M2 markers *Arg1* (B) and *Chil3* (**C**), as well as arginase enzymatic activity (**D**) in IL-13 - treated mouse BMDM. The treatment with IL-13 was performed in the presence of FBS-lo (<100 kDa) fraction as a source of growth factors (see Methods).

(**E-H**) Comparisons of effects ABS, ABS-lo (<100 kDa), and ABS-hi (>100 kDa) serum fractions on the expression of M1 markers *NOS2* and *TNF* on mRNA levels (**E, F**) and the expression of NOS2 on protein level (**G, H**) in IFNγ and LPS – treated murine BMDM. A representative image is shown in (**G**, see Methods) and quantification is shown in (**H**). The treatment with IFNγ and LPS was performed in serum-free DMEM media (see Methods).

For (**B-F**), (H): n=3 biological replicas; *$p<0.05$, **, $p<0.01$, data represented as mean ± s.e.m., one-way ANOVA followed by Tukey's posthoc test; **B**: $p<0.0001$, $F (5, 12) = 24.1$; **C**: $p<0.0001$, $F (5, 12) = 462.7$; **D**: $p<0.0001$, $F (4, 13) = 154.8$; **E**: $p<0.0001$, $F (4, 10) = 127.1$; **F**: $p<0.0001$, $F (4, 10) = 742.2$; **H**: $p<0.0001$, $F (4, 10) = 34.4$.

Abbreviations: ABS, adult bovine serum; ABS-hi, high molecular weight (>100 kDa) fraction of ABS; ABS-lo, low molecular weight (<100 kDa) fraction of ABS; FBS, fetal bovine serum; FBS-lo, low molecular weight (<100 kDa) fraction of FBS; Untr., untreated.

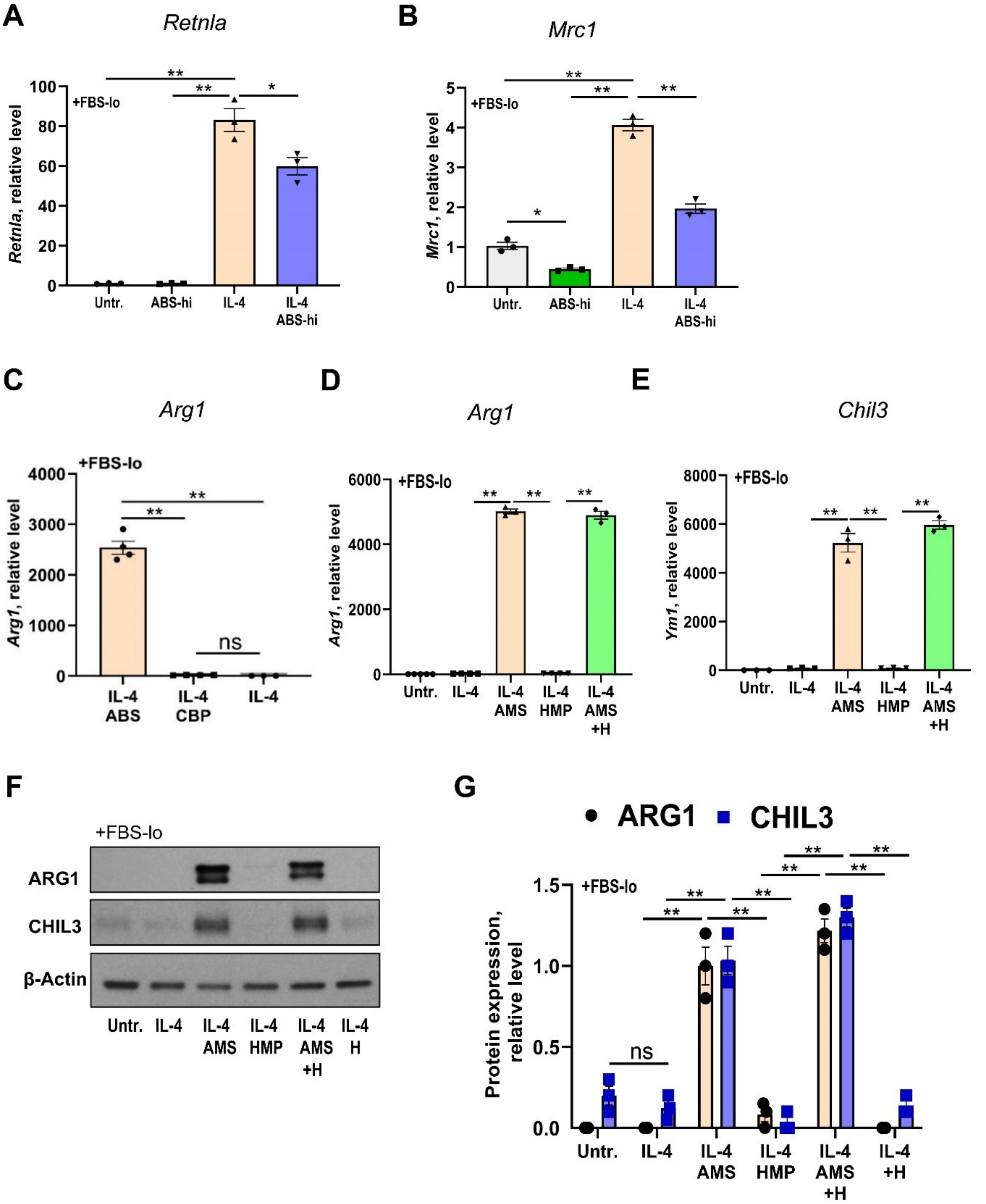


**<u>Figure S2</u>. Validation of the pattern of expression of IL-4-dependent genes and the influence of inhibition of blood coagulation on the functional activity of cofactor for IL-4 to induce the expression of M2 markers in macrophages.**

(**A, B**) Validation of the pattern of expressions of IL-4-dependent genes *Relma* (A) and *Mrc1* (B) by real-time RT PCR (see Methods and **Figure 2A**).

(**C**) Comparisons of adult bovine serum (ABS) with citrate bovine plasma (CBP) for their ability to induce the expression of Arg1 in mouse BMDM (see Methods and **Figure 3**).

(**D-G**) Comparisons of adult mouse serum (AMS), heparinized mouse plasma (HMP), and adult mouse serum with subsequently added heparin (AMS+H) on the expression of M2 markers Arg1 and Chil3 on mRNA (**D, E**) and protein (**F**, western blot image; **G**, quantification) levels in IL-4-treated vs. untreated murine BMDM (see **Figure 3**, Methods).

For (**A-E**), and (**G**): n=3 biological replicas; *$p<0.05$, **, $p<0.01$, data represented as mean ± s.e.m., one-way ANOVA followed by Tukey's posthoc test, **A**: $p<0.0001$, $F (3, 8) = 134.0$, **B**: $p<0.0001$, $F (3, 8) = 228.2$; **C**: $p<0.0001$, $F (2, 8) = 314.5$; **D**: $p<0.0001$, $F (4, 14) = 2870$; **E**: $p<0.0001$, $F (4, 10) = 260.7$; **G**, **ARG1**: $p<0.0001$, $F (5, 12) = 93.7$; **G, CHIL3**: $p<0.0001$, $F (5, 12) = 97.3$.

Abbreviations: ABS, adult bovine serum; AMS, adult mouse plasma; HMS, heparinized adult mouse plasma; H, heparin; FBS-lo, low molecular weight (<100 kDa) fraction of FBS; Untr., untreated.

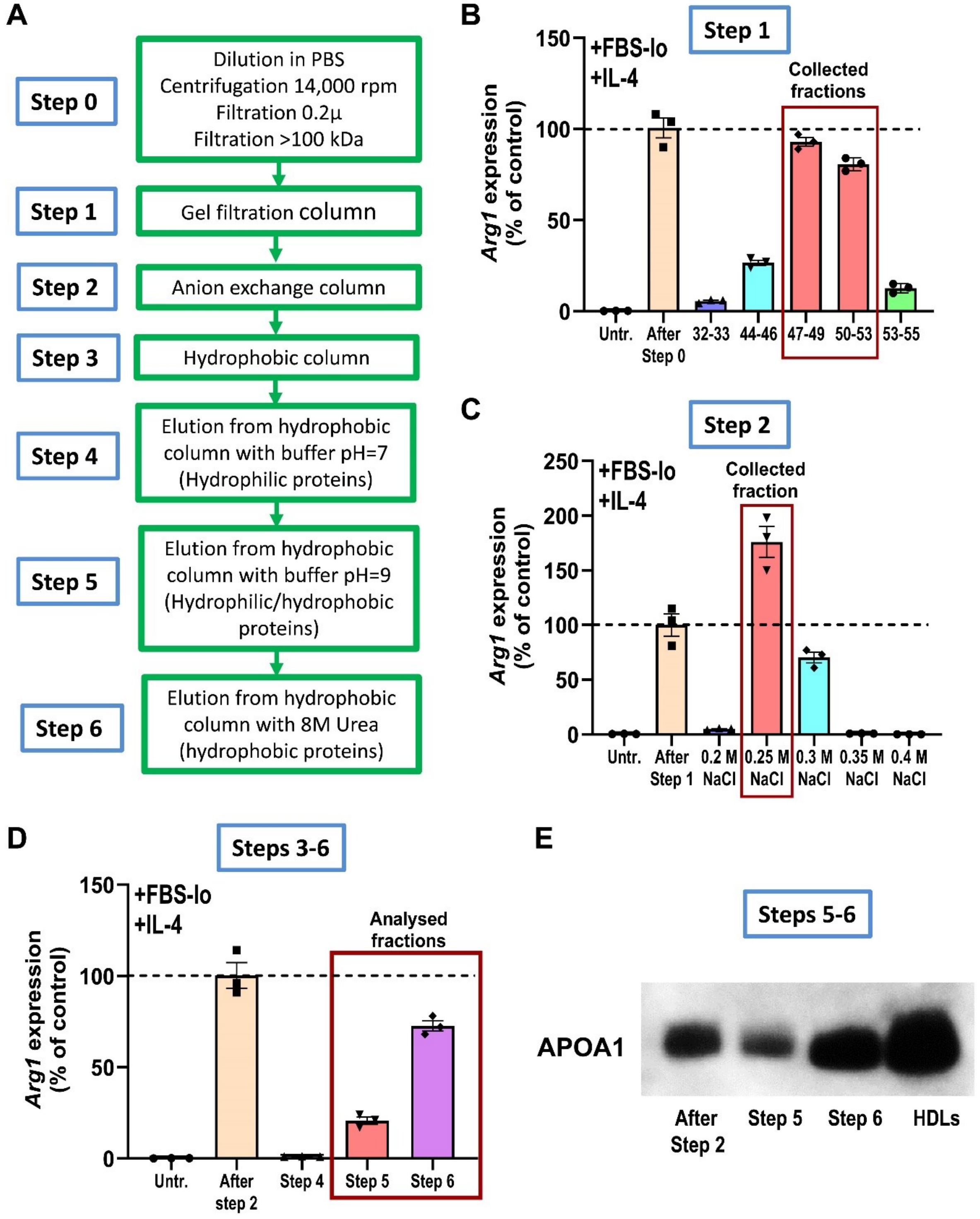


**<u>Figure S3</u>. Identification of HDLs in ABS-hi serum fraction as the most active proteins to induce *Arg1* expression in macrophages.**

(**A**) Experimental strategy for multistep protein fractionation of adult bovine serum using size exclusion (> 100 kDa) followed by gel filtration, anion exchange, and hydrophobic interaction columns (see Methods).

(**B-D**) Analysis of separated protein fractions for their biological activity to induce Arg1 expression in macrophages cell line after gel filtration (B, most active fractions 47-53) followed by anion exchange column (**C**, 0.25M NaCl active fraction) and the elution from hydrophobic column (**D**, Step 5 and Step 6 fractions; see Methods).

(**E**) Western blot analysis of APOA1 protein in the most biologically active fractions after Steps 5 and Step 6 fractionations compared to active fractions after the anion exchange column (after Step 2). Bovine HDLs were prepared by gradient ultracentrifugation (see Methods and **Figure 4A**) as the positive control. Loading protein concentrations were determined by Bradford assay (see Methods).

<u>Abbreviations</u>: FBS-lo, low molecular weight (<100 kDa) fraction of FBS; Untr., untreated.

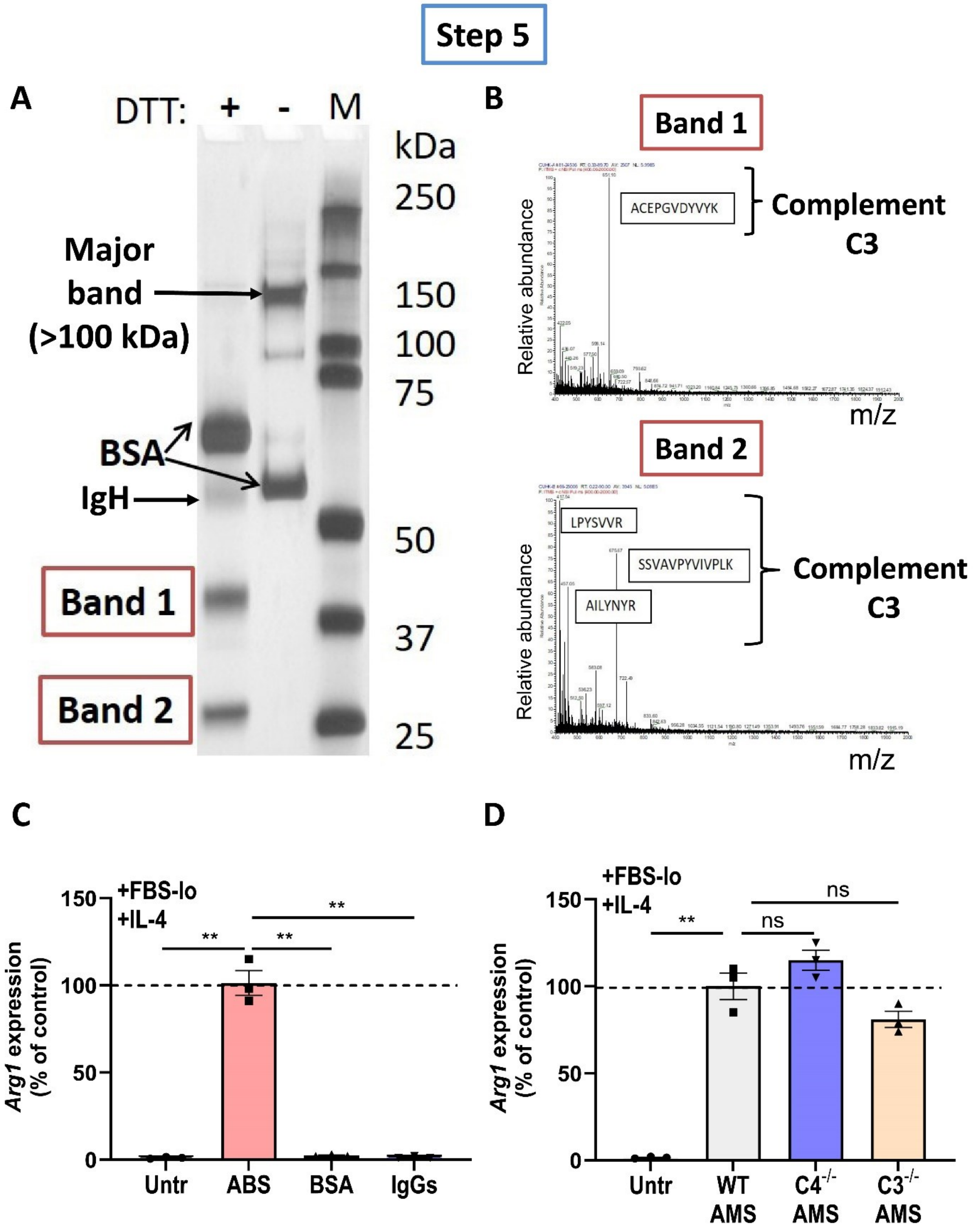


**<u>Figure S4</u>. Analysis of protein composition and biological activity of the protein fraction eluted from the hydrophobic column.**

(**A**) Gel image (silver staining for protein content) for analysis of heterogeneity of protein fraction after Step 5 (see **Figure S3**) during elution of proteins from the hydrophobic column with buffer pH=9 in the presence (left line) or absence of DTT (right line) treatment. With intact disulfide bonds (no DTT treatment), two major bands were detected that correspond to our band of interest with sizes in the range from 100 to 150 kDa and irrelevant BSA band with sizes in the range from 50 to 75 kDa (right line). After DDT treatment aimed to reduce disulfide bonds (see Methods), we still identified the presence of bovine serum albumin band (BSA, shifted slightly up after the addition of DDT), immunoglobulin heavy chain (IgH) band, and two unknown bands with the size in the range from 37 to 50 kDa (Band 1), and from 25 to 37 kDa (Band 2).

(**B**) In-gel proteinase digestion and LC-MS analysis of dissected unknown Band 1 and Band 2 revealed the presence of complement C3 as a major component of these bands and APOA1 and APOA4 as minor components (not shown).

(**C**) Comparisons of ABS, BSA, and bovine serum immunoglobulins G fraction (IgGs) for their ability to induce Arginase 1 expression in the macrophage cell line (see Methods).

(**D**) Comparisons of adult mouse serums (AMS) from wild-type (WT) vs. C3- or C4- deficient mice.

For (**C**), (**D**): **, $p<0.01$, ns, not significant, one-way ANOVA followed by Tukey's posthoc test; **C**: n=3 biological replicas, $p<0.0001$, $F (3, 8) = 194.9$; **D**: n=3 mice, $p<0.0001$, $F (3, 8) = 89.5$).

Abbreviations: ABS, adult bovine serum; BSA, bovine serum albumin; DTT, dithiothreitol; FBS-lo, low molecular weight (<100 kDa) fraction of FBS; IgGs, immunoglobulins of class G; IgH, heavy chain of immunoglobulin; Untr, untreated.

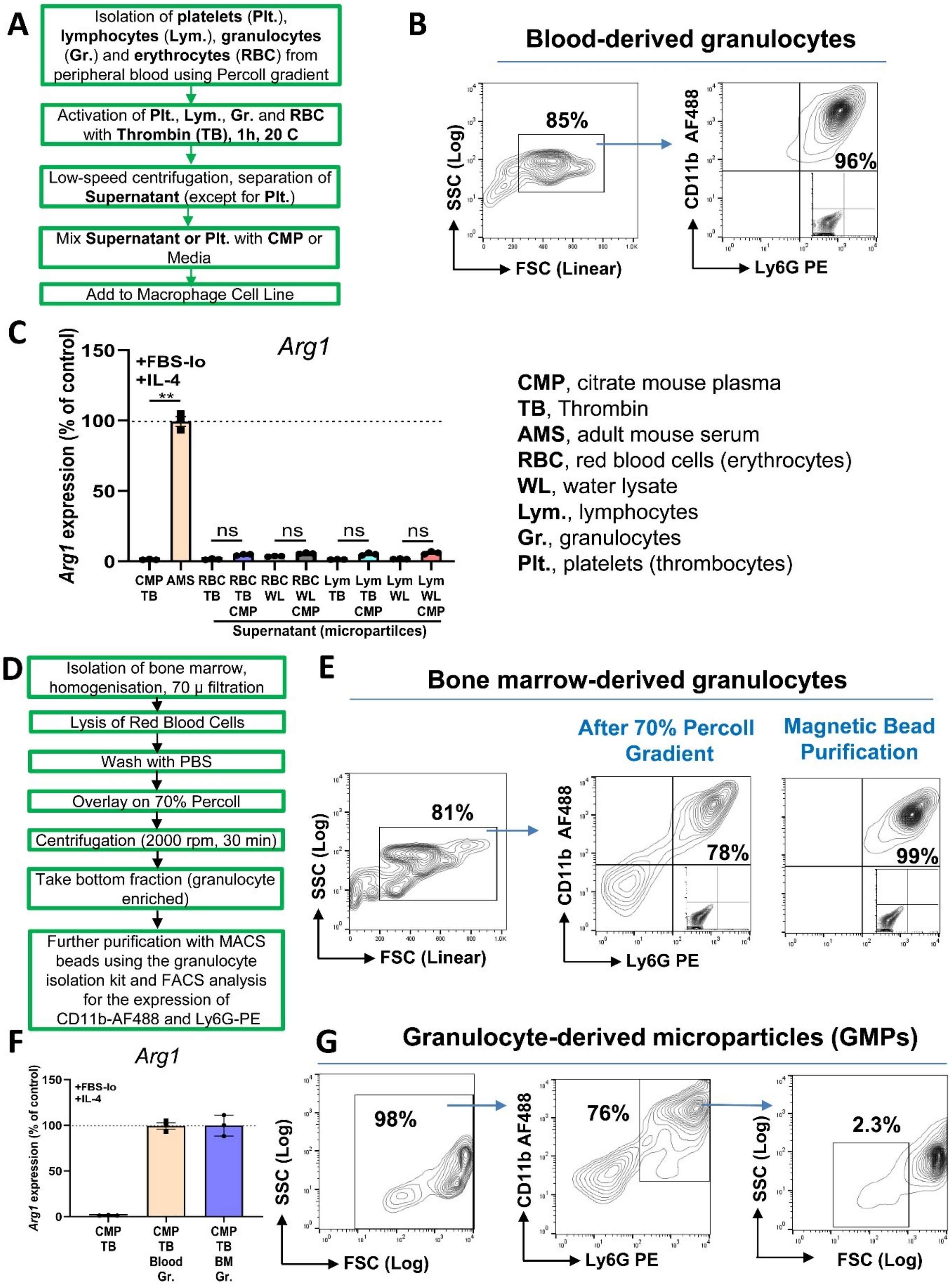


**Figure S5. Comparison of the ability of various blood-derived cell populations, blood-, and bone-marrow-derived granulocytes to induce Arg1 in macrophages.**
(**A**) Experimental design for isolation of platelets, mononuclear leukocytes ('lymphocytes', see Methods), granulocytes (neutrophils), and erythrocytes (red blood cells, RBCs).

(**B**) Gating strategy and flow cytometry analysis of purity of separated murine blood-derived granulocytes based on FSC/SSC parameters (live cells, left contour plot, see Methods) and fluorescent staining for granulocyte markers CD11b and Ly6G. For fluorescent marker staining, the quadrants were set based on negative control staining with isotype-matched control antibodies (shown in the low right quadrant of the right contour plot).

(**C**) Separated blood cell populations of RBCs and lymphocytes were activated with thrombin (TB) or lysed with water (water lysate, WL), and their supernatants after low-speed centrifugation were mixed with or without CMP and added to macrophages for analysis of *Arg1* expression (see and Methods).

(**D**) Experimental design for isolation of murine bone-marrow-derived granulocytes.

(**E**) Gating strategy and flow cytometry analysis of purity of isolated bone marrow-derived granulocytes with Percoll gradient and further purified with magnetic beads based on FSC/SSC parameters to separate debris and dead cells (left contour plot) and fluorescent staining for granulocytes markers CD11b and Ly6G (see Methods). For fluorescent marker staining, the quadrants were set based on negative control staining with isotype-matched control antibodies (shown in the low right quadrant of the right contour plot).

(**F**) Comparison of blood- vs. bone marrow-derived granulocytes to stimulate the expression of Arg1 in the macrophage cell line.

(**G**) Gating strategy of analysis of granulocyte-derived microparticles (GMPs) based on FSC/SSC parameters to the gate of alive cells and microparticles (left contour plot), and gating on $CD11b^{+}Ly6G^{+}$ cells (granulocytes and GMPs), and identification of FSC-low quadrant for identification of smaller size microparticles.

For (**C, F**): **, $p<0.01$; ns, not significant; n=3 biological replicas; one-way ANOVA followed by Tukey's posthoc test, **C**: $p<0.0001$, $F(9, 20) = 702.6$; **F**: $p<0.0001$, $F(2, 6) = 171.5$.

Abbreviations: see the Figure.

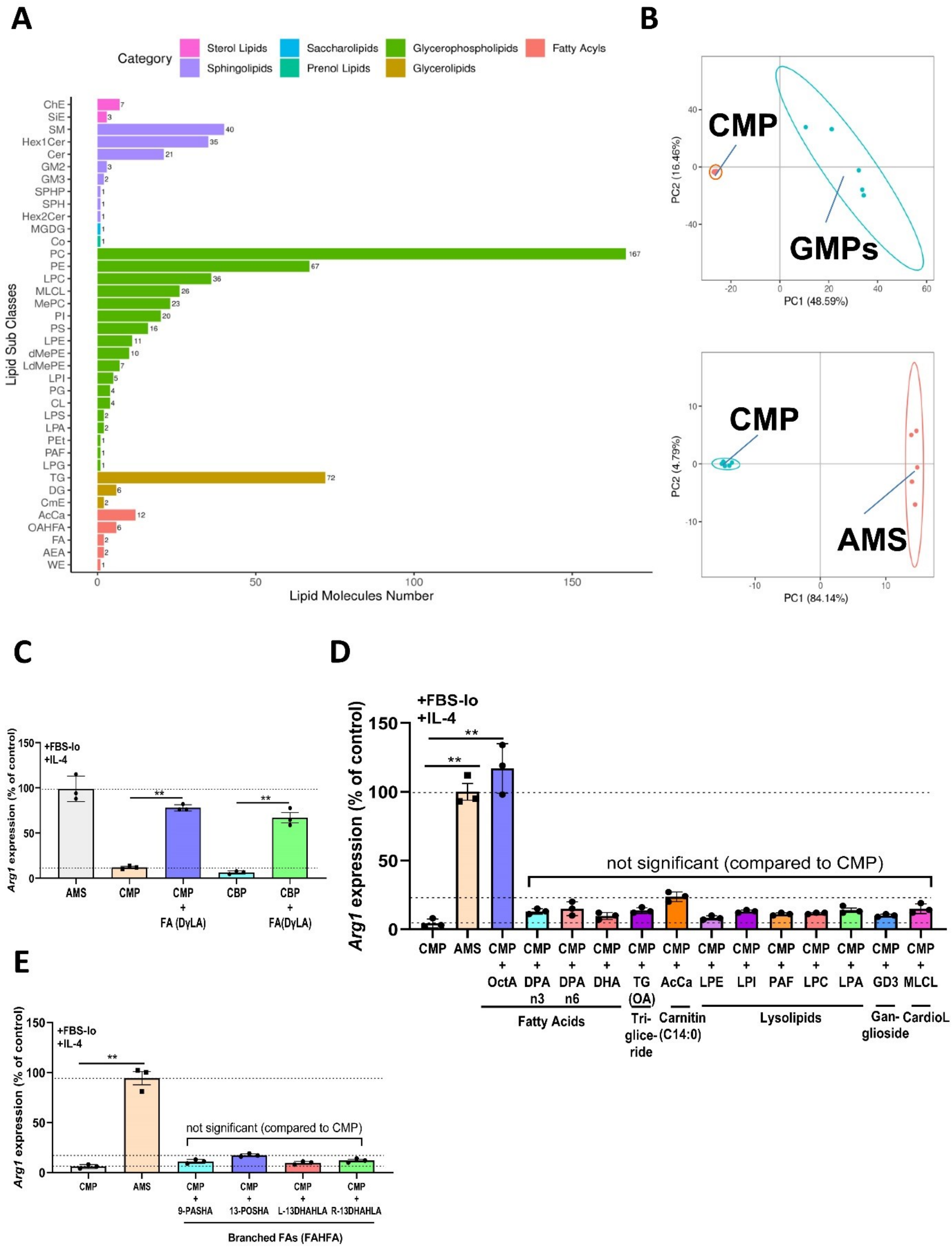


**Figure S6. Identification of subclasses of lipids by global untargeted lipidomics in CMP, AMS, and GMP samples, and the comparison of exogenous lipids to induce Arg1 expression in macrophages.**

(**A**) The vertical y-axis shows the lipid subclasses identified in the experiment, while the horizontal x-axis shows the number of identified lipid molecules in each lipid subclass. Different colors mark different lipid categories as shown on the top of the panel.

(**B**) Principal component analysis (PCA) demonstrated the separation of GMPs vs. CMP (left) and AMS vs. CMP (right) groups of five samples (for all: n=5 biological replicas).

(**C**) Functional analysis of adding exogenous fatty acid DγLA to CBP vs. CMP for their ability to induce *Arg1* expression in the macrophage cell line.

(**D, E**) Functional analysis of AMS- and granulocyte-derived lipids (lysolipids, triglycerides, carnitine, cardiolipin, sphingolipids, and non-branched fatty acids including DHA and DPA (**D**) and branched fatty acids (**E**)) for their ability to induce *Arg1* in the macrophage cell line.

For (**C-E**): **, $p<0.01$; n=3 biological replicas, one way ANOVA followed by Tukey's posthoc test; **C**: $p<0.0001$, $F(4, 10) = 81.2$; **D**: $p<0.0001$, $F(14, 30) = 100.4$; **E**: $p<0.0001$, $F(5, 12) = 138.9$).

Abbreviations: AMS, adult mouse serum; CBP, citrate bovine plasma; CMP, citrate mouse plasma; DHA, docosahexaenoic acid; DγLA, dihomo-γ-linolenic acid; DPA, docosapentaenoic acid; FAHFA, fatty acid esters of hydroxy fatty acids; GMPs, granulocyte-derived microparticles. PCA, principal component analysis.

A

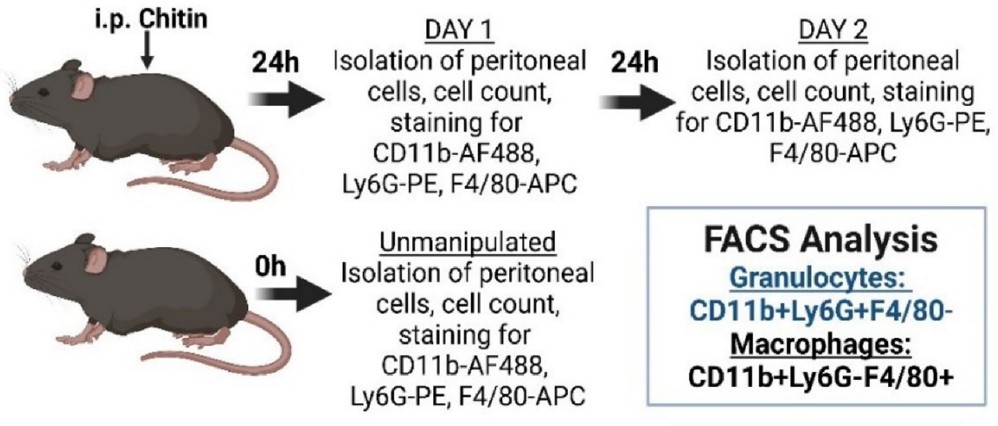


B

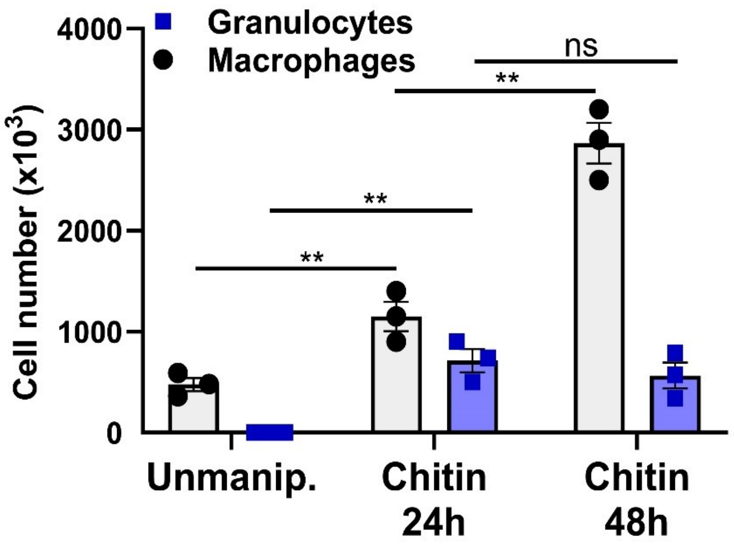


**<u>Figure S7.</u> Flow cytometry analysis of absolute numbers of macrophages and granulocytes during type 2 inflammation.**

(**A**, **B**) Experimental design (**A**) and flow cytometry (FACS) analysis (**B**) of absolute numbers of $CD11b^{+}F4/80^{+}Ly6G^{-}$ macrophages and $CD11b^{+}Ly6G^{+}F4/80^{-}$ granulocytes (see Methods) during type 2 inflammation 24-48 h after intraperitoneal injection of chitin.

For (**B**): **, $p<0.01$; ns, not significant; n=3 mice, one-way ANOVA followed by Tukey's posthoc test; Macrophages: $p<0.0001$, $F (2, 6) = 68.7$; Granulocytes: $p=0.0055$, $F (2, 6) = 14$.

<u>Abbreviations</u>: Unmanip., unmanipulated.

### SUPPLEMENTARY TABLES

**Table S1. Statistical Analysis for Figures 1-7**

| Figure (comparisons) | N | Test | P-value | F-value or T-value |
|---|---|---|---|---|
| **Fig.1A**, *Arg1* mRNA level (multiple comparisons) | 4 biological replicas (wells) | Ordinary one-way ANOVA followed by Tukey's posthoc test | P<0.0001 | F (5, 18) = 105.8 |
| **Fig.1B**, *Arg1* mRNA level (FBS+IL-4 vs. ABS+IL-4) | 4 biological replicas (wells) | Two-way ANOVA [Arg1 expression x time (days)] | P<0.0001 | F (1, 31) = 408.4 |
| **Fig.1B**, ARG1 protein level (FBS+IL-4 vs. ABS+IL-4) | 4 biological replicas (wells) | Two-way ANOVA [ARG1 expression x time (days)] | P<0.0001 | F (1, 20) = 332.2 |
| **Fig.1B** Arginase activity (multiple comparisons) | 4-6 biological replicas (wells) | Ordinary one-way ANOVA followed by Tukey's posthoc test | P<0.0001 | F (3, 16) = 282.8 |
| **Fig.1C**, FBS *Arg1* mRNA level (IL-4+FBS-lo vs. IL-4+FBS-hi) | 4 biological replicas (wells) | Ordinary one-way ANOVA followed by Tukey's posthoc test | P<0.0001 | F (2, 9) = 218.2 |
| **Fig.1C**, ABS *Arg1* mRNA level (IL-4+ABS-lo vs. IL-4+ABS-hi) | 4 biological replicas (wells) | Ordinary one-way ANOVA followed by Tukey's posthoc test | P<0.0001 | F (2, 9) = 618.2 |
| **Fig.1D**, *Arg1* mRNA level (multiple comparisons) | 3 biological replicas (wells) | Ordinary one-way ANOVA followed by Tukey's posthoc test | P<0.0001 | F (5, 12) = 276.1 |
| **Fig.1D**, *Chil3* mRNA level (multiple comparisons) | 3 biological replicas (wells) | Ordinary one-way ANOVA followed by Tukey's posthoc test | P<0.0001 | F (5, 12) = 754.4 |
| **Fig. 1F,** ARG1 protein level (multiple comparisons) | 3 biological replicas (wells) | Ordinary one-way ANOVA followed by Tukey's posthoc test | P<0.0001 | F (4, 10) = 92.8 |
| **Fig. 1F**, CHIL3 protein level (multiple comparisons) | 3 biological replicas (wells) | Ordinary one-way ANOVA followed by Tukey's posthoc test | P<0.0001 | F (4, 10) = 60.1 |
| **Fig. 1G,** Arginase activity (multiple comparisons) | 3 biological replicas (wells) | Ordinary one-way ANOVA followed by Tukey's posthoc test | P<0.0001 | F (4, 10) = 974.4 |

| | | | | |
|---|---|---|---|---|
| **Fig. 3B,** *Arg1* mRNA level (multiple comparisons) | 3 biological replicas (wells) | Ordinary one-way ANOVA followed by Tukey's posthoc test | P<0.0001 | F (4, 16) = 1972 |
| **Fig. 3B,** *Chil3,* mRNA level (multiple comparisons) | 3 biological replicas (wells) | Ordinary one-way ANOVA followed by Tukey's posthoc test | P<0.0001 | F (4, 10) = 33.7 |
| **Fig. 3D,** ARG1 protein level (multiple comparisons) | 3 biological replicas (wells) | Ordinary one-way ANOVA followed by Tukey's posthoc test | P<0.0001 | F (4, 10) = 194.8 |
| **Fig. 3D,** CHIL3 protein level (multiple comparisons) | 3 biological replicas (wells) | Ordinary one-way ANOVA followed by Tukey's posthoc test | P<0.0001 | F (4, 10) = 189.3 |
| **Fig. 3E,** Arginase activity (multiple comparisons) | 3 biological replicas (wells) | Ordinary one-way ANOVA followed by Tukey's posthoc test | P<0.0001 | F (4, 10) = 275.6 |
| **Fig. 3H,** OCR basal level (multiple comparisons) | 3 biological replicas (wells) | Ordinary one-way ANOVA followed by Tukey's posthoc test | P=0.0003 | F (3, 8) = 22.5 |
| **Fig. 3H,** OCR maximal level (multiple comparisons) | 3 biological replicas (wells) | Ordinary one-way ANOVA followed by Tukey's posthoc test | P<0.0001 | F (3, 8) = 467.3 |
| **Fig. 3I,** ECAR basal level (multiple comparisons) | 3 biological replicas (wells) | Ordinary one-way ANOVA followed by Tukey's posthoc test | P<0.0001 | F (3, 8) = 246.1 |
| **Fig. 3I,** ECAR maximal level (multiple comparisons) | 3 biological replicas (wells) | Ordinary one-way ANOVA followed by Tukey's posthoc test | P<0.0001 | F (3, 8) = 169.0 |
| **Fig. 4B**, ABS *Arg1* mRNA level (multiple comparisons) | 3 biological replicas (different ABS batches) | Ordinary one-way ANOVA followed by Tukey's posthoc test | P<0.0001 | F (4, 10) = 310.2 |
| **Fig. 4B**, AMS *Arg1* mRNA level (multiple comparisons) | 3 biological replicas (different AMS preparations) | Ordinary one-way ANOVA followed by Tukey's posthoc test | P<0.0001 | F (4, 10) = 34.5 |
| **Fig. 4D,** *Arg1* mRNA level | 3 biological replicas | Ordinary one-way ANOVA followed | P<0.0001 | F (6, 14) = 197.2 |

| | | | | |
|---|---|---|---|---|
| (multiple comparisons) | (different HDL preparations) | by Tukey's posthoc test | | |
| **Fig. 4F,** *Arg1* mRNA level (multiple comparisons) | 3 biological replicas (different cell preparations) | Ordinary one-way ANOVA followed by Tukey's posthoc test | P<0.0001 | F (7, 16) = 50.6 |
| **Fig. 5D,** % of GMPs (multiple comparisons) | 3 biological replicas (samples) | Ordinary one-way ANOVA followed by Tukey's posthoc test | P<0.0001 | F (3, 8) = 49.4 |
| **Fig. 5E,** TG release (multiple comparisons) | 3 biological replicas (samples) | Ordinary one-way ANOVA followed by Tukey's posthoc test | P<0.0001 | F (3, 8) = 66.5 |
| **Fig. 5F,** *Arg1* mRNA level (multiple comparisons) | 3 biological replicas (samples) | Ordinary one-way ANOVA followed by Tukey's posthoc test | P<0.0001 | F (4, 10) = 227.2 |
| **Fig. 5G,** *Arg1* mRNA level (multiple comparisons) | 3 mice (B6) | Ordinary one-way ANOVA followed by Tukey's posthoc test | P<0.0001 | F (7, 16) = 90.8 |
| **Fig. 6D**, C15:0 (AMS vs. CMP) | 3 biological replicas (different CMP and AMS preparations) | Unpaired Student's t-test | P=0.0004 | T (4) =-18.0 |
| **Fig. 6D**, C16:0 (AMS vs. CMP) | 3 biological replicas (different CMP and AMS preparations) | Unpaired Student's t-test | P<0.0001 | T (4) = -34.8 |
| **Fig. 6D**, C18:0 (AMS vs. CMP) | 3 biological replicas (different CMP and AMS preparations) | Unpaired Student's t-test | p=0.0002 | T (4) = -12.7 |
| **Fig. 6E**, C18:1 cis Linoleic Acid (LA) (AMS vs. CMP) | 3 biological replicas (different CMP and AMS preparations) | Unpaired Student's t-test | P<0.0001 | T (4) = -17.6 |
| **Fig. 6E**, C18:2 cis (AMS vs. CMP) | 3 biological replicas (different CMP and AMS preparations) | Unpaired Student's t-test | P<0.0001 | T (4) = -24.2 |
| **Fig. 6E**, C18:3 n3 (AMS vs. CMP) | 3 biological replicas (different CMP | Unpaired Student's t-test | P=0.0002 | T (4) = -13.0 |

| | | | | |
|---|---|---|---|---|
| | and AMS preparations) | | | |
| **Fig. 6E**, C18:3 n6 (AMS vs. CMP) | 3 biological replicas (different CMP and AMS preparations) | Unpaired Student's t-test | P=0.0003 | T (4) = -11.9 |
| **Fig. 6E**, C20:3 n6 (AMS vs. CMP) | 3 biological replicas (different CMP and AMS preparations) | Unpaired Student's t-test | P=0.0002 | T (4) = -13.4 |
| **Fig. 6E**, C20:4 n6 (AMS vs. CMP) | 3 biological replicas (different CMP and AMS preparations) | Unpaired Student's t-test | P<0.0001 | T (4) = -29.4 |
| **Fig. 6F**, *Arg1* mRNA level (CMP vs. AMS, or CMP vs. FAs) | 3 biological replicas (culture wells) | Ordinary one-way ANOVA followed by Tukey's posthoc test | P=0.0001 | F (12, 26) = 49.7 |
| **Fig. 6H,** *Arg1* mRNA level (multiple comparisons) | 3 biological replicas (culture wells) | Ordinary one-way ANOVA followed by Tukey's posthoc test | P<0.0001 | F (6, 14) = 93 |
| **Fig. 7A** Left panel. *Arg1* mRNA level (anti-SRB1 vs. anti-SRB3) | 3 biological replicas (culture wells) | Two-way ANOVA [*Arg1* expression x IgG concentration] | P<0.0001 | F (1, 20) = 71.0 |
| **Fig. 7A** Right panel. *Arg1* mRNA level (multiple comparisons) | 3 biological replicas (culture wells) | Ordinary one-way ANOVA followed by Tukey's posthoc test | P<0.0001 | F (4, 10) = 289.8 |
| **Fig 7B**, *Arg1* mRNA level (multiple comparisons) | 3 mice (B6) | Ordinary one-way ANOVA followed by Tukey's posthoc test | P<0.0001 | F (5, 12) = 247.2 |
| **Fig 7B**, *Chil3* mRNA level (multiple comparisons) | 3 mice (B6) | Ordinary one-way ANOVA followed by Tukey's posthoc test | P<0.0001 | F (5, 12) = 168.4 |
| **Fig 7C**, *Arg1* mRNA level (multiple comparisons) | 3 mice (B6) | Ordinary one-way ANOVA followed by Tukey's posthoc test | P<0.0001 | F (2, 6) = 603.4 |
| **Fig 7C**, *Chil3* mRNA level | 3 mice (B6) | Ordinary one-way ANOVA followed | P<0.0001 | F (2, 6) = 128.1 |

| | | | | |
|---|---|---|---|---|
| (multiple comparisons) | | by Tukey's posthoc test | | |
| **Fig 7D**, *Arg1* mRNA level (PBS vs. AMS) | 3 mice (B6) | Unpaired Student's t-test | P<0.0001 | T (4) = 18.4 |
| **Fig 7D**, *Chil3* mRNA level (PBS vs. AMS) | 3 mice (B6) | Unpaired Student's t-test | P=0.0001 | T (4) = 15.24 |

**<u>Table S2.</u> Primer Sequences for Gene Expression Analysis**

| Gene | Primer | Sequence |
|---|---|---|
| ***GADPH*** | Forward | 5'-ATGACCACAGTCCATGCCATC-3' |
| | Reverse | 5'-GAGCTTCCCGTTCAGCTCTG-3' |
| ***Arg1*** | Forward | 5'CTTGGCTTGCTTCGGAACTC-3' |
| | Reverse | 5'- GGAGAAGGCGTTTGCTTAGTTC-3' |
| ***Chil3 (Ym1)*** | Forward | 5'-CCATTGGAGGATGGAAGTTTG-3' |
| | Reverse | 5'- GACCCAGGGTACTGCCAGTC-3' |
| ***Retnla (Relmα)*** | Forward | 5'-GCCAGGTCCTGGAACCTTTC-3' |
| | Reverse | 5'-GGAGCAGGGAGATGCAGATGAG-3' |
| ***NOS2*** | Forward | 5'-ACCCACATCTGGCAGAATGAG-3' |
| | Reverse | 5'-AGCCATGACCTTTCGCATTAG-3' |
| ***TNF*** | Forward | 5'-AGCCGATGGGTTGTACCTTG-3' |
| | Reverse | 5'- GTGGGTGAGGAGCACGTAGTC-3' |
